# Autocrine STAT3 activation in HPV positive cervical cancer through a virus-driven Akt - NFκB - IL-6 signalling axis

**DOI:** 10.1101/507954

**Authors:** Ethan L. Morgan, Andrew Macdonald

## Abstract

Persistent human papillomavirus (HPV) infection is the leading cause of cervical cancer. Although the fundamental link between HPV infection and oncogenesis is established, the specific mechanisms of virus-mediated transformation remain poorly understood. We previously demonstrated that the HPV encoded E6 protein increases the activity of the proto-oncogenic transcription factor STAT3 in primary human keratinocytes; however, the molecular basis for STAT3 activation in cervical cancer remains unclear. Here, we show that STAT3 phosphorylation in HPV positive cervical cancer cells is mediated primarily via autocrine activation by the pro-inflammatory cytokine Interleukin 6 (IL-6). Antibody-mediated blockade of IL-6 signalling in HPV positive cells inhibits STAT3 phosphorylation, whereas both recombinant IL-6 and conditioned media from HPV positive cells leads to increased STAT3 phosphorylation within HPV negative cervical cancer cells. Interestingly, we demonstrate that non-conventional activation of the transcription factor NFκB, involving the protein kinase Akt, is required for IL-6 production and subsequent STAT3 activation. Our data provides new insights into the molecular re-wiring of cancer cells by HPV E6. We reveal that activation of an IL-6 signalling axis drives the autocrine and paracrine phosphorylation of STAT3 within HPV positive cervical cancers cells. Greater understanding of this pathway provides a potential opportunity for the use of existing clinically approved drugs for the treatment of HPV-mediated cervical cancer.

**Author Summary:** Persistent infection with HPV is the predominant cause of anogenital and oral cancers. Transformation requires the re-wiring of signalling pathways in infected cells by virus encoded oncoproteins. At this point a comprehensive understanding of the full range of host pathways necessary for HPV-mediated carcinogenesis is still lacking. In this study we describe a signalling circuit resulting in the aberrant production of the IL-6 cytokine. Mediated by the HPV E6 oncoprotein, it requires activation of the NFκB transcription factor. The autocrine and paracrine actions of IL-6 are essential for STAT3 activation in HPV-positive cervical cancers. This study provides molecular insights into the mechanisms by which a virus encoded oncoprotein activates an oncogenic pathway, and illuminates potential targets for therapeutic intervention.

## Introduction

Human papillomaviruses (HPV) are a leading cause of squamous cell carcinomas of the ano-genital and oropharyngeal epithelium [1]. High risk HPVs (HR-HPV), exemplified by HPV16 and 18, are responsible for >99% of cervical, and between 30 – 70% of oropharyngeal cancers [2]. HPV encodes three oncogenic proteins: E5, E6 and E7, which manipulate signalling pathways necessary for cellular transformation. These include: epidermal growth factor receptor (EGFR) [3-5], Wnt [6, 7] and Hippo signalling [8]. The E5 membrane protein activates EGFR signalling [9] by a mechanism linked to its virus-coded ion channel (viroporin) activity [3, 10, 11]. HPV E6 forms complexes with host E3 ubiquitin ligases and mediates proteasomal degradation of the p53 tumour suppressor protein, as well as increasing telomerase activity in order to prevent apoptosis and immortalise infected cells [12]. The E7 oncoprotein stimulates the DNA damage response, driving viral replication and genomic instability, simultaneously promoting the progression of cells through the S phase of the cell cycle [13, 14].

The transcription factor signal transducer and activator of transcription (STAT) 3 is an essential regulator of cellular proliferation, differentiation and survival [15]. It is a *bona fide* oncogene and its aberrant activation has been observed in a growing number of malignancies [16]. As such, STAT3 has become an attractive therapeutic target in a diverse range of cancers, including bladder, ovarian and head and neck squamous cell carcinoma (HNSCC) [17].

Oncogenic viruses can activate STAT3 to drive cellular proliferation, necessary for viral replication and tumourigenesis [18]. Using a primary keratinocyte cell culture model, we previously demonstrated that E6 activates STAT3 signalling during the productive HPV lifecycle [19]. STAT3 activation was essential for the hyperplasia observed in HPV-containing keratinocyte raft culture models. Increased STAT3 protein expression and phosphorylation also correlated with cervical disease progression in a panel of cytology samples [19]. Although we identified that Janus kinase 2 (JAK2) and MAP kinases were necessary for STAT3 phosphorylation in HPV-containing primary keratinocytes, our understanding of the mechanisms by which E6 mediates this process remains incomplete. Furthermore, the mechanisms underpinning STAT3 activation in cervical cancer cells also lacks a comprehensive understanding, whereas inhibition of STAT3 activity results in a profound reduction in cellular proliferation and the induction of apoptosis [20, 21].

A number of extracellular stimuli including cytokines and growth factors induce STAT3 phosphorylation and signalling [17]. This requires the phosphorylation of tyrosine 705 (Y705) and serine 727 (S727), resulting in STAT3 dimerisation and nuclear translocation, where it is able to regulate gene expression [16]. In particular, members of the IL-6 family of cytokines are key mediators of STAT3 activation through their interactions with the gp130 co-receptor [22].

Here, we show that HPV positive cervical cancer cells have higher levels of phosphorylated STAT3 protein when compared with those that are HPV negative. This results from increased IL-6 production and release, leading to autocrine and paracrine activation of STAT3 via a signalling pathway requiring the IL-6 co-receptor gp130. Additionally, we show that IL-6 production is controlled by E6-mediated stimulation of NF-κB signalling, which appears to be partially dependent on a non-canonical activation of the pathway requiring the Akt protein kinase. Finally, we demonstrate a correlation between IL-6 expression and cervical disease progression, suggesting that targeting the IL-6 pathway to prevent STAT3 activation may have therapeutic benefits in cervical cancer.

## Results

### STAT3 protein expression and phosphorylation is increased in HPV positive compared with HPV negative cervical cancer cells

To establish the benchmark for HPV-mediated augmentation of STAT3 phosphorylation, we first analysed the level of STAT3 phosphorylation in a panel of six cervical cancer cell lines. These included two HPV negative (HPV-) lines (C33A and DoTc2), two HPV16 positive (HPV16+; SiHa and CaSKi) and two HPV18 positive (HPV18+; SW756 and HeLa) lines. Both HPV16+ and HPV18+ cancer cells displayed a higher level of STAT3 phosphorylation at both (Y705 and S727) sites compared to the HPV- cell lines (significant in SiHa, CaSKi and HeLa, p<0.05). Additionally, the overall abundance of STAT3 was increased in the HPV+ cervical cancer cells compared with HPV- cervical cancer cells (Fig 1A and B). Together, these data demonstrate increased STAT3 expression and phosphorylation within HPV+, compared with virus negative, cervical cancer cells.

**Figure 1.**
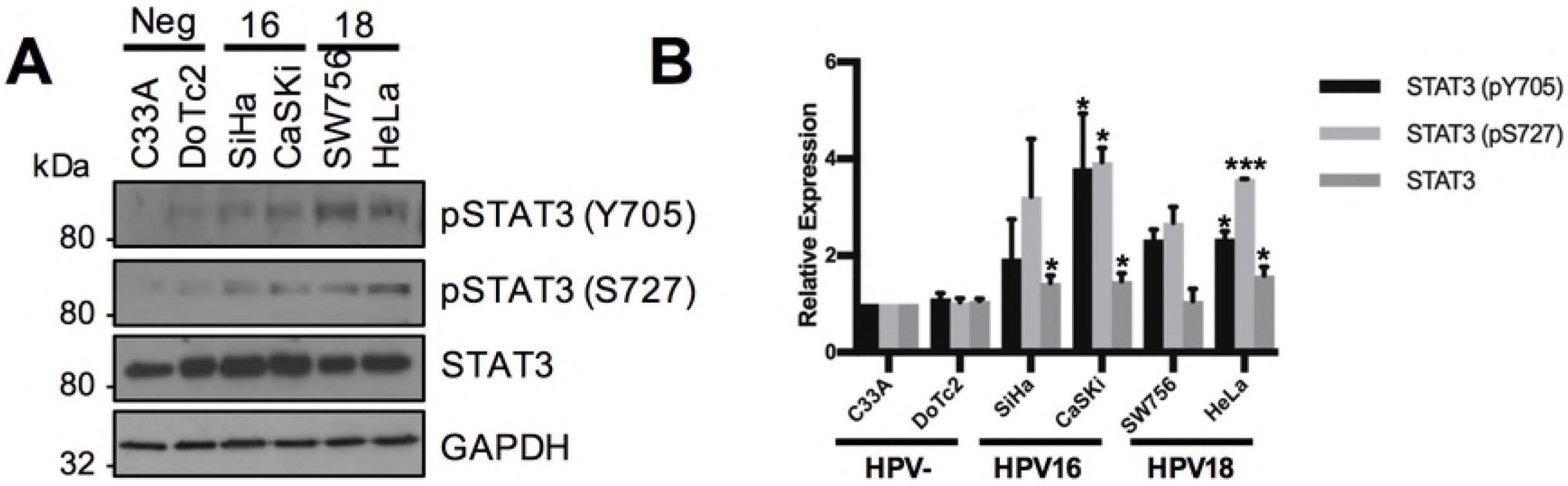
STAT3 phosphorylation is higher in HPV+ verses HPV- cervical cancer cells. **A)** Representative western blot of from six cervical cancer cell lines – two HPV-(C33A and Dotc2 4510), two HPV16+ (SiHa and CaSKi) and HPV18+ (SW756 and HeLa) – for the expression of phosphorylated (Y705 and S727) and total STAT3. GAPDH served as a loading control. Data are representative of at least three biological independent repeats. **B)** Quantification of the protein band intensities in **A)** standardised to GAPDH levels. Bars represent the means ± standard deviation from at 3 independent biological repeats. *P<0.05 (Student’s t-test).

### A secreted factor in the media of HPV+ cells can induce STAT3 phosphorylation in HPV- cervical cancer cells

Given that STAT3 is often regulated by extracellular signals, we investigated whether HPV promotes the secretion of factors capable of inducing STAT3 phosphorylation. C33A cells (HPV-) incubated with conditioned media (CM) from HeLa cells (HPV18+) displayed marked STAT3 phosphorylation on both Y705 and S727 residues, reaching a peak between 30 minutes and 1 hour (Fig 2A). Similar results were observed with CaSKi-CM (HPV16+) (Fig 2B). Accordingly, HeLa or CaSKi-CM induced increased STAT3 nuclear accumulation within C33A cells (Fig 2C and D). These data indicate that a secreted factor in the media from HPV+ cells induces STAT3 phosphorylation in target cells.

**Figure 2.**
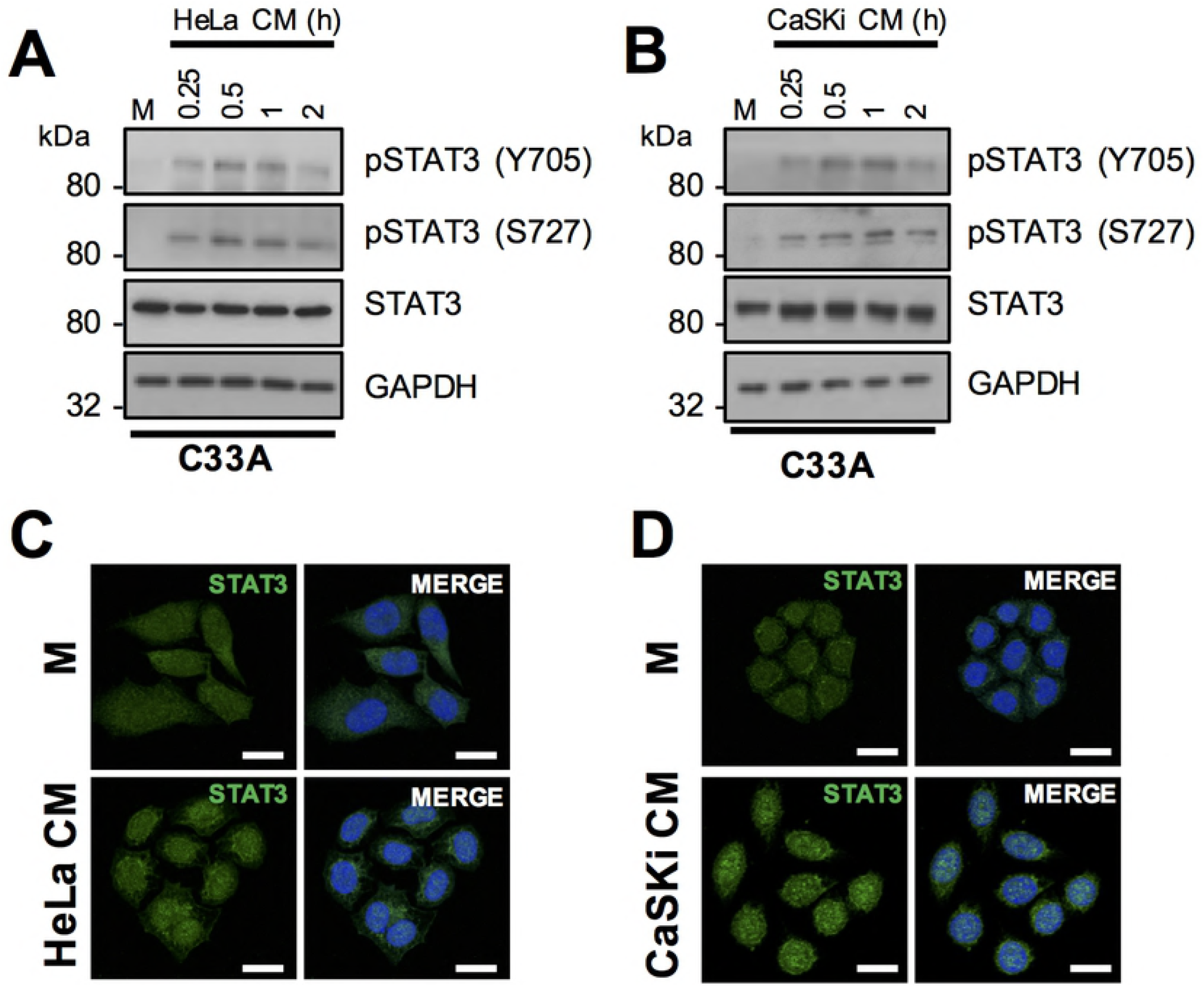
A secreted factor from HPV+ cervical cancer cells can induce STAT3 phosphorylation in HPV- cervical cancer cells. **A-B)** C33A cells were serum starved for 24 hours and conditioned media from **A)** HeLa or **B)** CaSKi cells was added for the indicated time points. For the control, C33A conditioned media was added to cells for 2 hours (M in the figure). Represented western blot shows cell lysates analysed for the expression of phosphorylated and total STAT3. GAPDH served as a loading control. Data are representative of at least three biological independent repeats. **C-D)** C33A cells were serum starved for 24 hours and incubated with conditioned media from **C)** HeLa or **D)** CaSKi cells for 2 hours. For the control, C33A conditioned media was added to cells for 2 hours (M in the figure). Cells were analysed by immunofluorescence staining for total STAT3 (green) and counterstained with DAPI to highlight the nuclei (blue in the merged panels). Scale bar, 20 μm.

### IL-6 secretion is increased in HPV+ cervical cancer cells

To identify the secreted factor responsible for inducing STAT3 phosphorylation, we focused on members of the IL-6 family of pro-inflammatory cytokines, as these have a well-studied role in the activation of STAT3 [23]. Firstly, the mRNA expression levels of key members of the family were analysed by RT-qPCR. In both HeLa and CaSKi cells, *IL6, IL10, LIF* (Leukaemia inhibitory factor) and *OSM* (Oncostatin M) mRNA levels were significantly higher than in C33A cells (Fig 3A), with *IL-6* showing the greatest increase. Building on this, we analysed *IL-6* mRNA expression in all six cervical cancer cell lines. In both HPV16+ and HPV18+ cervical cancer cells, a significantly higher level of *IL-6* mRNA expression was observed compared with HPV- cervical cancer cells (Fig 3B), which correlated with intracellular IL-6 protein expression when analysed by western blot (Fig 3C). Finally, to confirm that the IL-6 protein detected was secreted, an IL-6 specific ELISA was performed. IL-6 was undetectable in the culture medium of HPV- cells; in contrast a significant quantity of IL-6 could be detected the culture medium of both HPV16+ and HPV18+ cells (SiHa, p = 0.03; CaSKi, p = 0.0004; SW756, p = 0.03; HeLa, p = 0.00004). These data suggest that IL-6 expression and secretion is much higher in HPV+ cervical cancer cells compared with uninfected lines.

**Figure 3.**
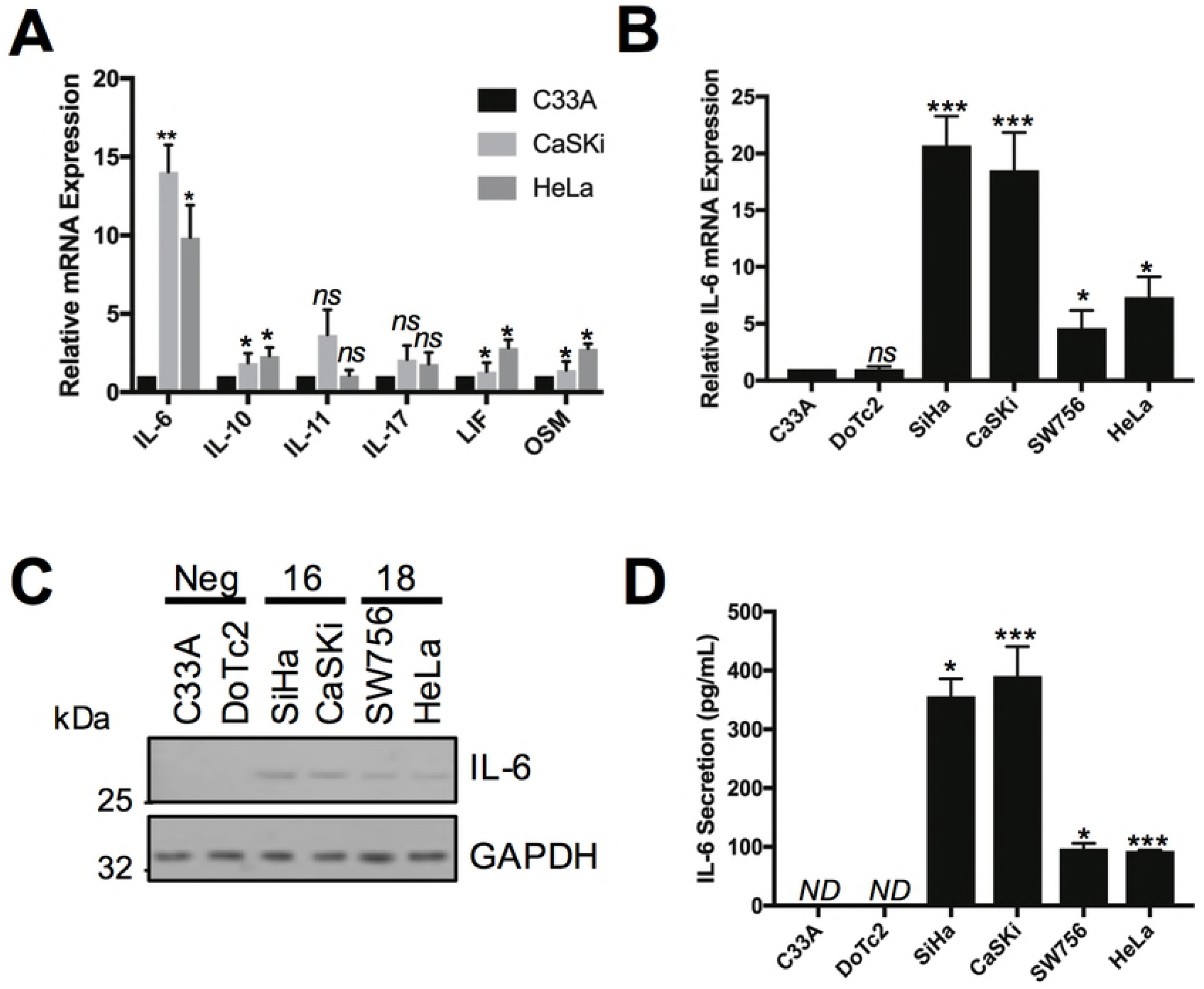
Interleukin-6 (IL-6) is up regulated in HPV+ cervical cancer cells. **A)** The expression level of cytokines from the IL-6 family were analysed in HPV-, HPV16+ and HPV18+ cervical cancer cells by qRT-PCR. Samples were normalized against U6 mRNA levels. Representative data are presented relative to the HPV- cervical cancer cells. Bars are the means ± standard deviation from at least three biological repeats. *P<0.05, **P<0.01, ***P<0.001 (Student’s t-test). **B)** The expression of IL-6 from six cervical cancer cell lines – two HPV- (C33A and Dotc2 4510), two HPV16+ (SiHa and CaSKi) and HPV18+ (SW756 and HeLa) – was analysed by qRT-PCR. Samples were normalized against U6 mRNA levels. Representative data are presented relative to the HPV- cervical cancer cells. Bars are the means ± standard deviation from at least three biological repeats. *P<0.05, **P<0.01, ***P<0.001 (Student’s t-test). **C)** Representative western blot of from six cervical cancer cell lines – two HPV- (C33A and Dotc2 4510), two HPV16+ (SiHa and CaSKi) and HPV18+ (SW756 and HeLa) – for the expression of IL-6. GAPDH served as a loading control. Data are representative of at least three biological independent repeats. **D)** ELISA analysis from the culture medium from six cervical cancer cell lines – two HPV- (C33A and Dotc2 4510), two HPV16+ (SiHa and CaSKi) and HPV18+ (SW756 and HeLa) – for secreted IL-6 protein. Error bars represent the mean +/- standard deviation of a minimum of three biological repeats. ND = not determined (below the detection threshold). *P<0.05, **P<0.01, ***P<0.001 (Student’s t-test).

### IL-6 binding to gp130-containing receptor complexes is required for STAT3 phosphorylation and nuclear translocation in cervical cancer cells

IL-6 signalling is initiated by an interaction between IL-6 and the IL-6 receptor (IL-6R) – gp130 co-receptor complex [17]. In cervical cancer cells, blocking antibodies against IL-6 and gp130 were utilised to confirm that they were required for STAT3 phosphorylation. Firstly, we confirmed that IL-6 was the mediator of STAT3 phosphorylation secreted by HPV+ cells by pre-incubating C33A cells with the gp130 blocking antibody before treating them with HeLa or CaSKi-CM. Separately, we added the neutralising IL-6 antibody to the CM before addition to cells. CM from HPV+ cells induced STAT3 phosphorylation in HPV- cells; however, pre-incubation of C33A cells with gp130 blocking antibody, or the addition of CM containing the neutralising IL-6 antibody, blocked the ability of HPV+ conditioned media to induce STAT3 phosphorylation (Fig 4A) and nuclear translocation (Fig 4B). This suggests that IL-6 secretion from HPV+ cervical cancer cells is able to induce the paracrine activation of STAT3 in HPV- cervical cancer cells via IL-6/gp130 signalling.

**Figure 4.**
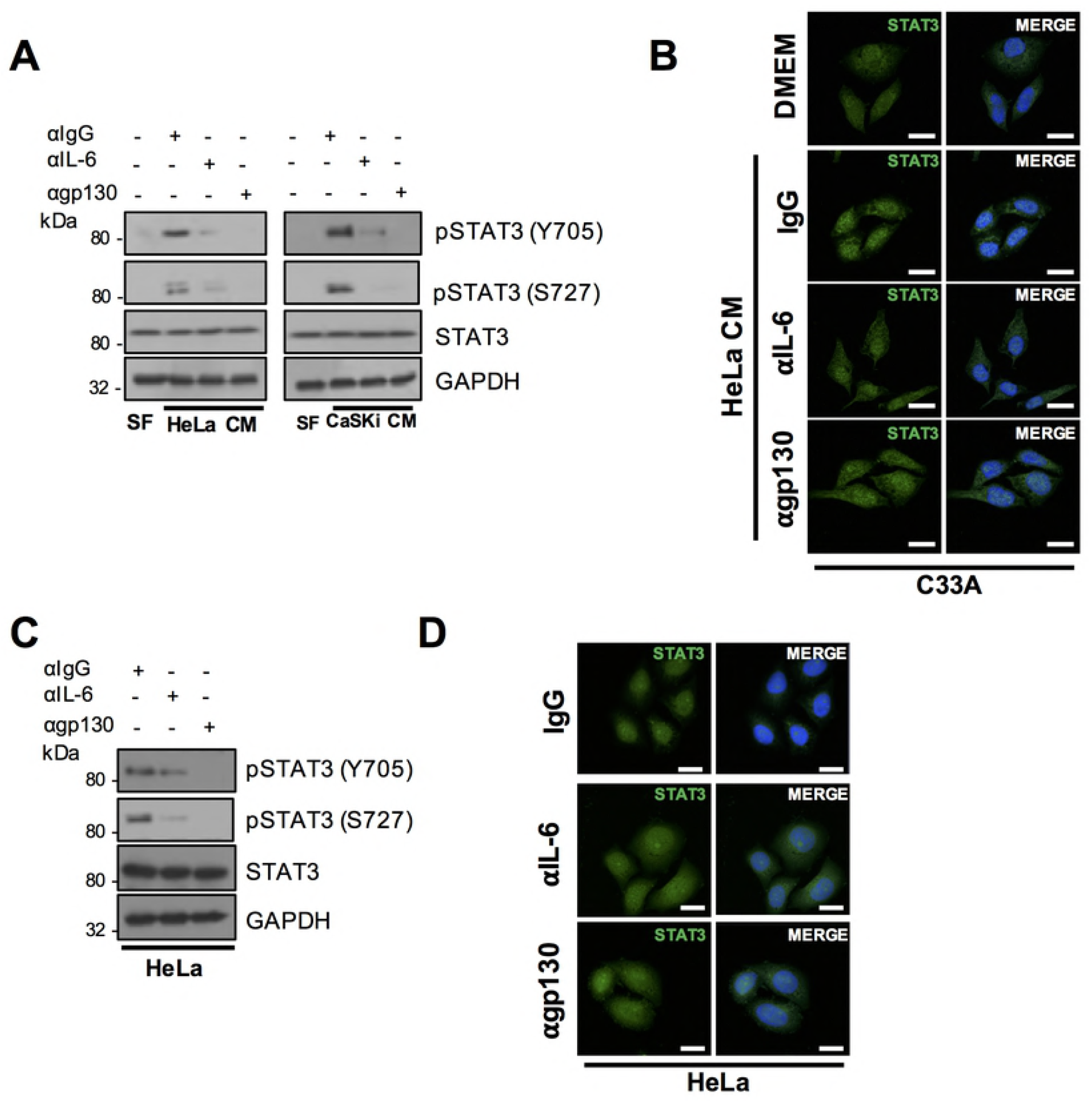
IL-6/gp130 signalling is required for STAT3 phosphorylation and nuclear translocation in cervical cancer cells. **A)** Representative western blot of C33A cells treated with conditioned medium (CM) from HeLa and CaSKi cells for 2 hours. Cells were pre-treated with IgG, anti-IL6 or anti-gp130 antibody for 4 hours before CM addition. Cell lysates were analysed for phosphorylated and total STAT3 expression. GAPDH served as a loading control. Data are representative of at least three biological independent repeats. **B)** C33A cells treated with conditioned medium (CM) from HeLa and CaSKi cells for 2 hours. Cells were pre-treated with IgG, anti-IL6 or anti-gp130 antibody for 4 hours before CM addition. Cells were then analysed by immunofluorescence staining for total STAT3 (green) and counterstained with DAPI to highlight the nuclei (blue in the merged panels). Scale bar, 20 μm. **C)** HeLa cells were treated with IgG, anti-IL6 or anti-gp130 for 4 hours. Cell lysates were analysed for phosphorylated and total STAT3 expression. GAPDH served as a loading control. Data are representative of at least three biological independent repeats. **D)** HeLa cells were treated with IgG, anti-IL6 or anti-gp130 for 4 hours. Cells were then analysed by immunofluorescence staining for total STAT3 (green) and counterstained with DAPI to highlight the nuclei (blue in the merged panels). Scale bar, 20 μm.

Finally, to confirm that IL-6/gp130 signalling was required for the autocrine activation of STAT3 in HPV+ cervical cancer cells, HeLa cells were pre-incubated with IL-6 and g130 neutralizing antibodies. Incubation with either neutralising antibody led to a reduction in STAT3 phosphorylation on both phosphorylation sites, accompanied by a block in STAT3 nuclear translocation, suggesting that IL-6 is required for the autocrine activation of STAT3 (Fig 4C and D). Taken together, these data demonstrate that HPV induces autocrine and paracrine IL-6/STAT3 signalling in cervical cancer.

### HPV E6 induction of IL-6 expression is required for STAT3 phosphorylation

The increased STAT3 phosphorylation observed in HPV containing keratinocytes is dependent on the E6 oncoprotein [19]. Additionally, HPV16 E6 has been previously demonstrated to induce IL-6 secretion in non-small cell lung cancer (NSCLC) cells [24]. Therefore, the ability of E6 to induce IL-6 expression in cervical cancer cells was assessed. To this end, we first transfected C33A with an IL-6 promoter luciferase reporter in combination with a GFP-E6 expression plasmid or the GFP vector. Expression of HPV18 E6 significantly increased IL-6 promoter activity by ~4-fold compared with the GFP control (Fig 5A; p= 0.002). This corresponded to a ~4-fold increase in endogenous *IL-6* mRNA expression (Fig 5B; p= 0.01) and IL-6 protein expression (Fig 5C). Finally, E6 expression resulted in a significant increase in IL-6 secretion (Fig 5D; p= 0.0245). To demonstrate that endogenous E6 could induce IL-6 expression in HPV+ cells, HeLa (Fig 5) or CaSKi (Supplementary Fig 1) cells were treated with two E6 specific siRNAs. Knockdown of E6 expression led to a significant reduction in *IL-6* mRNA expression (Fig 5E; p= 0.046 and Supplementary Fig 1A), IL-6 protein expression (Fig 5F and Supplementary Fig 1B) and secretion (Fig 5G; p= 0.003 and Supplementary Fig 1C). Together, these data demonstrate that HPV E6 increases the expression and secretion of IL-6.

**Figure 5.**
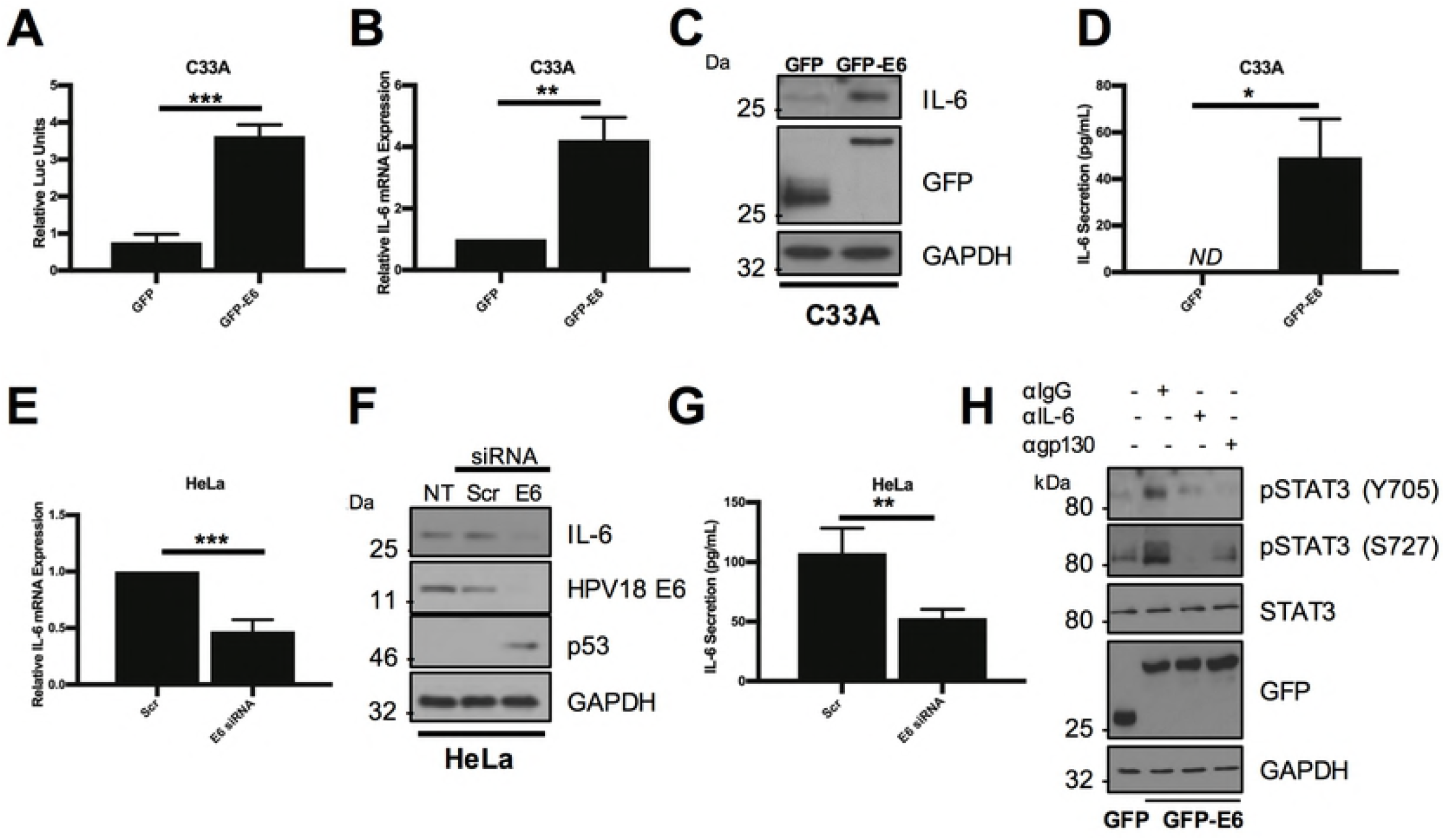
HPV E6 induced IL-6 expression is required for STAT3 phosphorylation. **A)** Representative luciferase reporter assay from C33A cells co-transfected with GFP tagged E6 and an IL-6 promoter reporter. Promoter activity was measured using a dual-luciferase system. Data are presented as relative to the GFP transfected control. **B)** C33A cells were transiently transfected with GFP or GFP tagged HPV18 E6 and RNA was extracted for qRT-PCR analysis of IL-6 expression. Samples were normalized against U6 mRNA levels. Representative data are presented relative to the GFP control. **C)** Representative western blot of C33A cells transiently transfected with GFP or GFP tagged HPV18 E6 and analysed for IL-6 expression. Expression of HPV E6 was confirmed using a GFP antibody and GAPDH served as a loading control. **D)** C33A cells were transiently transfected with GFP or GFP tagged HPV18 E6. The culture medium was analysed for IL-6 protein by ELISA. **E)** HeLa cells were transfected with a pool of two specific siRNAs against HPV18 E6 and analysed for IL-6 mRNA expression by qRT-PCR. Samples were normalized against U6 mRNA levels. **F)** Representative western blot of HeLa cells transfected with a pool of two specific siRNAs against HPV18 E6 and analysed for the expression of IL-6. Knockdown of HPV18 E6 was confirmed using an antibody against HPV18 E6 and p53. GAPDH served as a loading control. **G)** HeLa cells were transfected with a pool of two specific siRNAs against HPV18 E6. The culture medium was analysed for IL-6 protein by ELISA. **H)** Representative western blot of C33A cells transiently transfected with GFP or GFP tagged HPV18 E6 and treated with IgG, anti-IL6 or anti-gp130 for 4 hours before harvest. Cell lysates were then analysed for phosphorylated and total STAT3. Expression of HPV E6 was confirmed using a GFP antibody and GAPDH served as a loading control. Bars represent the means ± standard deviation from at least three independent biological repeats. *P<0.05, **P<0.01 (Student’s t-test).

To confirm if IL-6 expression was necessary for E6-mediated STAT3 phosphorylation, HPV18 E6 was expressed in C33A cells which were then treated with neutralising antibodies against either IL-6 or the gp130 co-receptor. As we have previously shown, the expression of HPV E6 induced STAT3 phosphorylation at both Y705 and S727 residues in C33A cells [19]. Treatment of E6 expressing cells with neutralising antibodies against IL-6 or gp130 led to the loss of STAT3 phosphorylation (Fig 5H). This suggests that the E6-mediated induction of IL-6 expression is essential for the autocrine phosphorylation of STAT3 in HPV+ cervical cancer cells.

### HPV E6 activates NFκB to induce IL-6 expression

Several signalling pathways can induce IL-6 expression, including the transcription factor NFκB, which is activated in response to a range of extracellular ligands such as TNFα [25]. HPV E6 has previously been shown to activate NFκB signalling under hypoxic conditions [26, 27]. To assess whether NFκB is necessary for the increased IL-6 expression, we first tested whether expression of E6 in isolation would activate NFκB in C33A cells. Using an NFκB driven luciferase reporter plasmid, overexpression of E6 induced NFκB activity ~3-fold compared to a GFP control (Fig 6A; p= 0.001).

**Figure 6.**
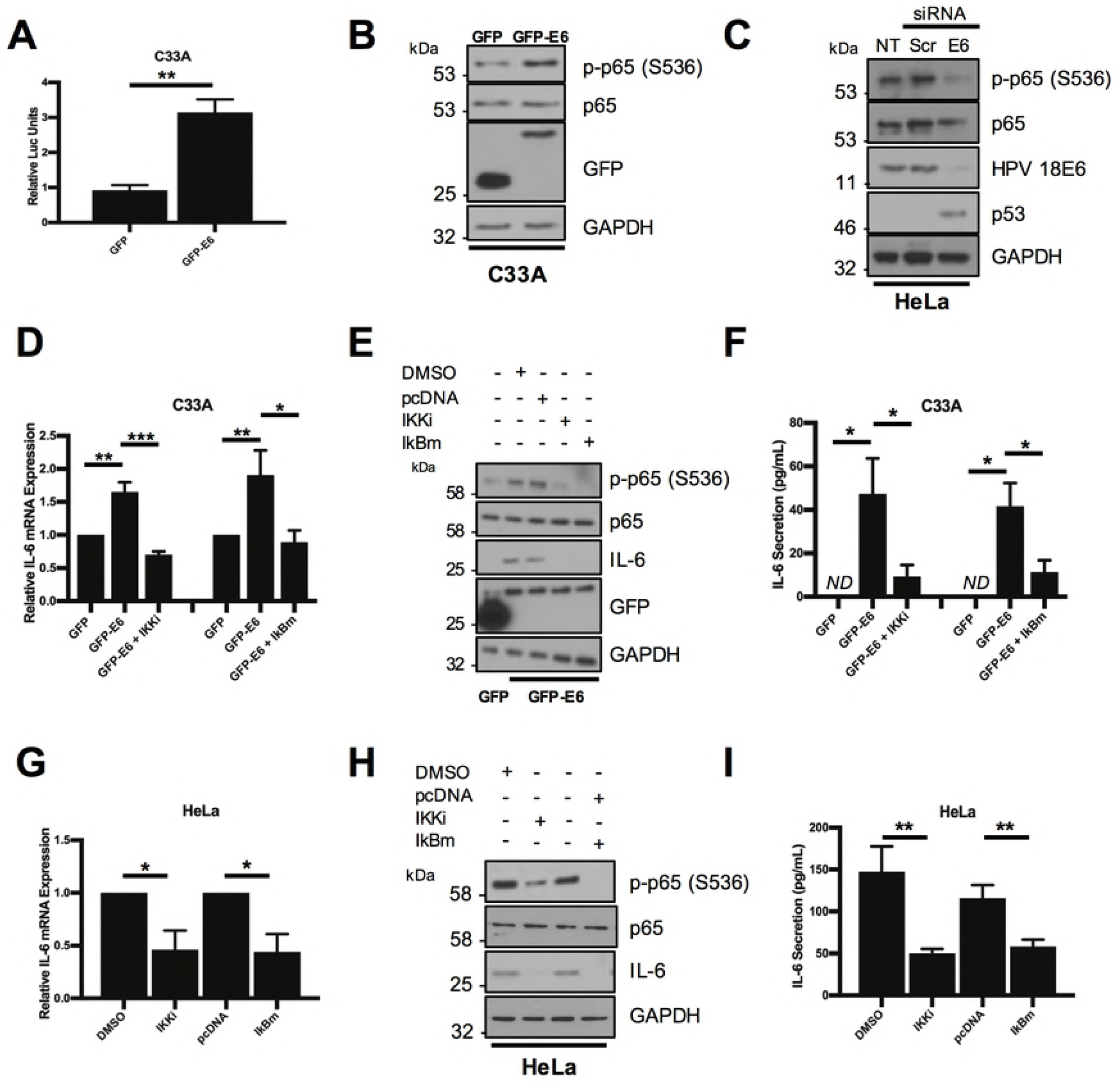
HPV E6 mediated IL-6 expression requires NF-κB activity. **A)** Representative luciferase reporter assay from C33A cells co-transfected with GFP tagged E6 and a ConA reporter containing tandem κB binding sites. Promoter activity was measured using a dual-luciferase system. Data are presented as relative to the GFP transfected control. **B)** Representative western blot of C33A cells transiently transfected with GFP or GFP tagged HPV18 E6 and analysed for phosphorylated and total p65 expression. Expression of HPV E6 was confirmed using a GFP antibody and GAPDH served as a loading control. **C)** Representative western blot of HeLa cells transfected with a pool of two specific siRNAs against HPV18 E6 and analysed for the expression of phosphorylated and total p65. Knockdown of HPV18 E6 was confirmed using an antibody against HPV18 E6 and p53. GAPDH served as a loading control. **D-F)** C33A cells were co-transfected with GFP, GFP tagged HPV18 E6 or GFP tagged HPV18 E6 and mutant IκBα (IκBm). Cells were then either left untreated or treated with IKK inhibitor VII (IKKi). **D)** Total RNA was extracted for qRT-PCR analysis of IL-6 expression. Samples were normalized against U6 mRNA levels. Representative data are presented relative to the GFP control. **E)** Cell lysates were analysed for the expression of phosphorylated and total p65 and IL-6. Expression of HPV E6 was confirmed using a GFP antibody and GAPDH served as a loading control. **F)** The culture medium was analysed for IL-6 protein by ELISA. **G-I)** HeLa cells transfected with a pool of two specific siRNAs against HPV18 E6. **G)** Total RNA was extracted for qRT-PCR analysis of IL-6 expression. Samples were normalized against U6 mRNA levels. Representative data are presented relative to the GFP control. **H)** Cell lysates were analysed for the expression of phosphorylated and total p65 and IL-6. Expression of HPV E6 was confirmed using a GFP antibody and GAPDH served as a loading control. **I)** The culture medium was analysed for IL-6 protein by ELISA. Bars represent the means ± standard deviation from at least three independent biological repeats. *P<0.05, **P<0.01, ***P<0.001 (Student’s t-test).

Canonical NFκB signalling results in the phosphorylation of the p65 subunit and its nuclear translocation, where it is transcriptionally active in complex with additional NFκB subunits including p50 [25]. Over expression of E6 in C33A cells induced robust p65 phosphorylation, without affecting total p65 protein levels (Fig 6B). In contrast, siRNA knockdown of E6 in HeLa cells reduced p65 phosphorylation (Fig 6C), together suggesting that HPV E6 activates NFκB signalling.

To assess if NFκB activity was required for the increase in IL-6 production observed in E6 expressing cells, we employed a dual approach to prevent NFκB activation in C33A cells overexpressing E6. Cells were treated either with a small molecule inhibitor (IKKi) targeting the IKKα/β complex, which is required to phosphorylate and activate NFκB or transfected with a plasmid encoding a mutant IκBα protein (IκBm), which cannot be degraded and as such retains NFκB in the cytosol in an inactive form [28]. Expression of E6 increased *IL-6* mRNA production, and inhibition of NFκB using either IKKi or IκBm led to a significant reduction in *IL-6* mRNA expression (Fig 6D; IKKi, p=0.00001; IκBm, p=0.04), IL-6 protein levels (Fig 6E) and secretion (Fig 6F; IKKi, p=0.02; IκBm, p=0.007). Importantly, both strategies effectively inhibited NFκB activity as judged by a reduction in p65 phosphorylation (Fig 6E).

To confirm if NFκB activity was also required for mediating the increased IL-6 levels seen in HPV+ cancer cells, NFκB activity was blocked in HeLa cells. Inhibition of NFκB led to a reduction in *IL-6* mRNA expression (Fig 6F; IKKi, p= 0.03; IκBm, p =0.02), IL-6 protein levels (Fig 6H) and secretion (Fig 6I; IKKi, p= 0.007; IκBm, p=0.007). Collectively, these data demonstrate that HPV E6-mediated IL-6 expression is dependent on NFκB activity.

### NFκB is required for STAT3 phosphorylation in E6 expressing cells

As NFκB was required for the increase in IL-6 expression observed in HPV+ cancer cell lines, it was necessary to next test whether NFκB was also required for the activation of STAT3. First, we tested the ability of an inducer of NFκB activation, TNFα, to induce STAT3 phosphorylation. Treatment of serum starved C33A cells with TNFα increased p65 phosphorylation, which peaked at 0.5 hours after treatment (Fig 7A; lane 3). TNFα treatment also induced IL-6 expression; starting at 0.5 hours post treatment and remained high up to 24 hours post treatment. TNFα treatment also led to an increase in STAT3 phosphorylation at both Y705 and S727 residues; however, this peaked later, at 2 hours (Fig 7A; lane 5). TNFα-dependent nuclear translocation of STAT could also be observed 2 hours post treatment (Fig 7B).

**Figure 7.**
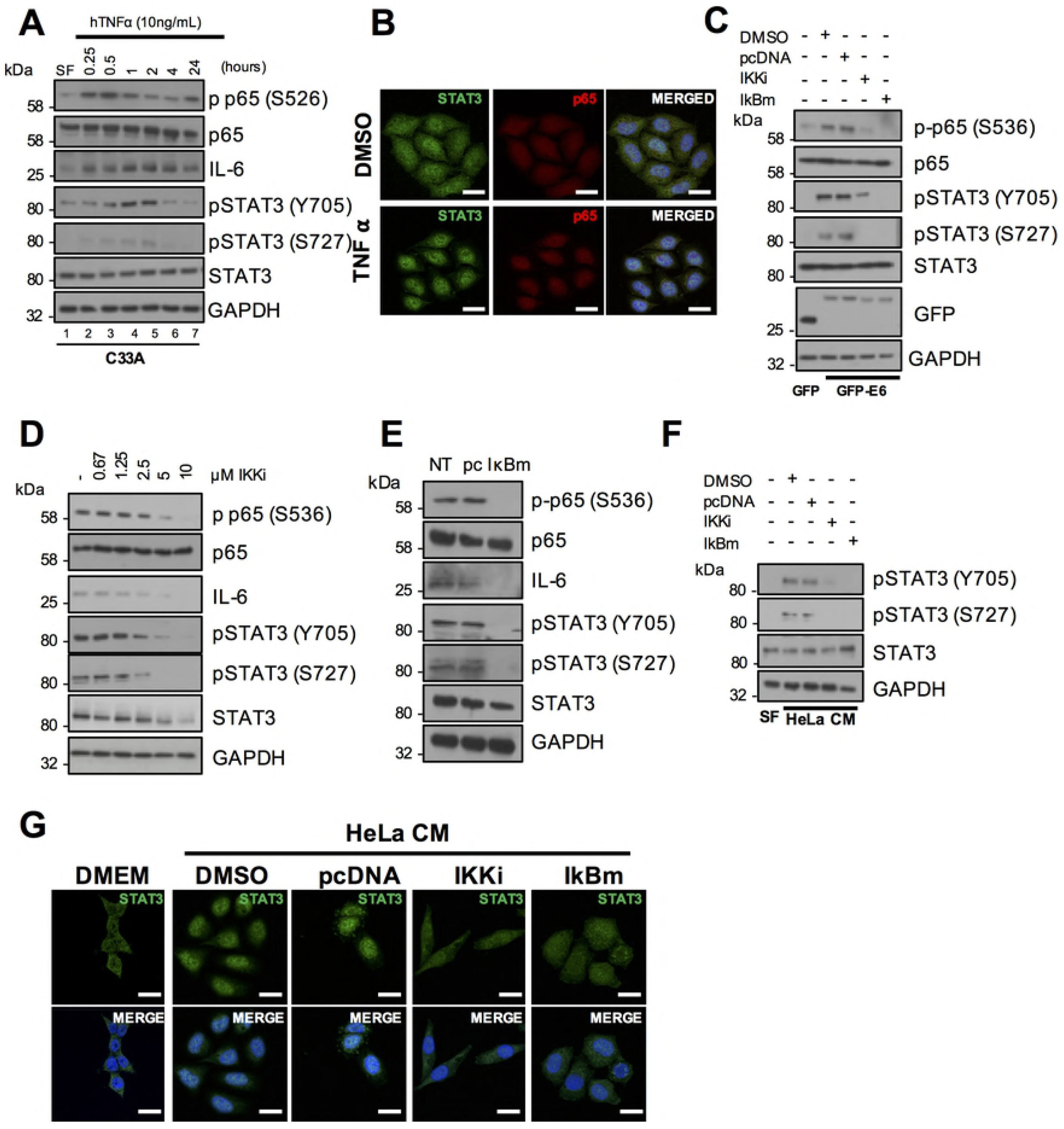
NF-κB activity is required for HPV E6 mediated STAT3 signalling. **A)** Representative western blot of C33A cells treated with 20 ng/mL recombinant human TNFα for the indicated time points. Cell lysates were analysed for phosphorylated and total p65, phosphorylated and total STAT3 and IL-6 expression. GAPDH served as a loading control. Data are representative of at least three biological independent repeats. **B)** C33A cells treated with 20 ng/mL recombinant human TNFα for 60 mins were fixed and were analysed by immunofluorescence staining for total STAT3 (green) and total p65 (red) and counterstained with DAPI to highlight the nuclei (blue in the merged panels). Scale bar, 20 μm. **C)** C33A cells were co-transfected with GFP, GFP tagged HPV18 E6 or GFP tagged HPV18 E6 and mutant IκBα (IκBm). Cells were then either left untreated or treated with IKK inhibitor VII (IKKi). Cell lysates were then analysed for phosphorylated and total p65, phosphorylated and total STAT3 expression. Expression of HPV E6 was confirmed using a GFP antibody and GAPDH served as a loading control. **D)** Representative western blot from HeLa cells treated with increasing doses of the IKKα/β inhibitor IKK inhibitor VII (IKKi). Cell lysates were analysed for the expression of phosphorylated and total p65, phosphorylated and total STAT3 and IL-6 expression. GAPDH served as a loading control. **E)** Representative western blot from HeLa cells transfected with mutant IκBα (IκBm). Cell lysates were analysed as in **D). F)** C33A cells were serum starved for 24 hours. Cells were then treated with HeLa condition media from HeLa cells treated with DMSO or IKKi or transfected with pcDNA or IκBm. Cell lysates were analysed for phosphorylated and total STAT3 expression. GAPDH served as a loading control. **G)** C33A cells were serum starved for 24 hours. Cells were then treated with HeLa condition media from HeLa cells treated with DMSO or IKKi or transfected with pcDNA or IκBm. Cells were analysed by immunofluorescence staining for total STAT3 (green) and counterstained with DAPI to highlight the nuclei (blue in the merged panels). Scale bar, 20 μm.

To assess the importance of NFκB activation for STAT3 phosphorylation, HPV E6 was first overexpressed in C33A cells, with or without treatment with the NFκB inhibitor IKKi, or co-expression of IκBm. E6 overexpression noticeably increased p65 and STAT3 phosphorylation and inhibition of NFκB using either approach led to a loss of STAT3 phosphorylation (Fig 7C), Additionally, blockade of NFκB also led to a reduction in STAT3 phosphorylation in HeLa cells (Fig 7D and E), suggesting that E6 mediated STAT3 phosphorylation is depended on NFκB activity. Finally, to ascertain if NFκB was essential for the paracrine activation of STAT3 in C33A cells, we took conditioned media from HeLa cells in which NFκB was inhibited and added this to C33A cells. This conditioned media failed to induce STAT3 phosphorylation (Fig 7F) and nuclear translocation (Fig 7G). Importantly, inhibition of NFκB activity had no effect on STAT3 phosphorylation or nuclear translocation mediated by treatment with exogenous IL-6 (Supp Fig 2A and 2B), demonstrating that NFκB is upstream of IL-6 secretion. Together, these data suggest that NFκB is required for the autocrine and paracrine activation of STAT3.

### The protein kinase Akt contributes to NFκB activation and IL-6 production in E6 expressing cells

NFκB is activated by multiple upstream signalling components in a stimulus and tissue-dependent manner [29]. The PI3K/Akt signalling pathway is frequently activated in cervical cancers due to mutations in the *PIK3CA* gene [30], and Akt can activate NFκB in some cancers [31, 32]. Furthermore, Akt is a known mediator of IL-6 expression [33]. We therefore assessed if Akt was involved in NFκB activation and IL-6 secretion in HPV+ cervical cancer. First, we confirmed that E6 can activate Akt, as measured by an increase in Akt phosphorylation, as has been previously shown [34]. Over expression of E6 in C33A cells led to a marked increase in Akt phosphorylation at both threonine 308 and serine 473, without affecting total Akt protein levels (Fig 8A). In addition, siRNA knockdown of E6 in HeLa cells reduced Akt phosphorylation (Fig 8B), together confirming that HPV E6 activates Akt.

**Figure 8.**
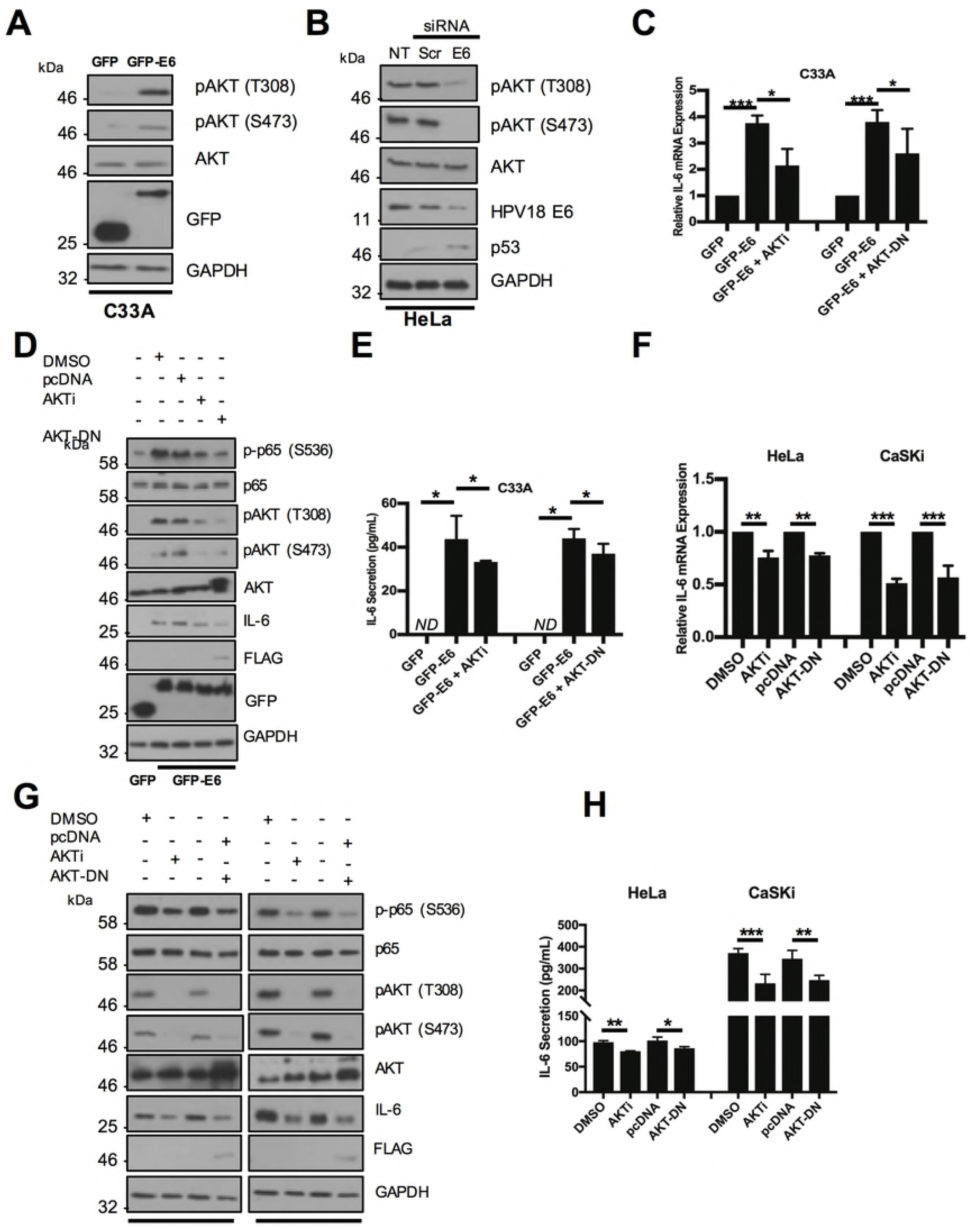
Activation of Akt by HPV E6 contributes to IL-6 expression via NF-κB. **A)** Representative western blot of C33A cells transiently transfected with GFP or GFP tagged HPV18 E6 and analysed for phosphorylated and total Akt expression. Expression of HPV E6 was confirmed using a GFP antibody and GAPDH served as a loading control. **B)** Representative western blot of HeLa cells transfected with a pool of two specific siRNAs against HPV18 E6 and analysed for the expression of phosphorylated and total Akt. Knockdown of HPV18 E6 was confirmed using an antibody against HPV18 E6 and p53. GAPDH served as a loading control. **C-E)** C33A cells were co-transfected with GFP, GFP tagged HPV18 E6 or GFP tagged HPV18 E6 and mutant Akt (Akt-DN). Cells were then either left untreated or treated with Akt inhibitor VIII (Akti). **C)** Total RNA was extracted for qRT-PCR analysis of IL-6 expression. Samples were normalized against U6 mRNA levels. Representative data are presented relative to the GFP control. **D)** Cell lysates were analysed for the expression of phosphorylated and total p65 and IL-6. Expression of HPV E6 was confirmed using a GFP antibody and GAPDH served as a loading control. **E)** The culture medium was analysed for IL-6 protein by ELISA **F-H)** HeLa and CaSki cells transfected with Akt-DN or treated with Akti. **F)** Total RNA was extracted for qRT-PCR analysis of IL-6 expression. Samples were normalized against U6 mRNA levels. Representative data are presented relative to the GFP control. **G)** Cell lysates were analysed for the expression of phosphorylated and total p65 and IL-6. Expression of HPV E6 was confirmed using a GFP antibody and GAPDH served as a loading control. **H)** The culture medium was analysed for IL-6 protein by ELISA. Bars represent the means ± standard deviation from at least three independent biological repeats. *P<0.05, **P<0.01, ***P<0.001 (Student’s t-test).

To interrogate the contribution of Akt activation to IL-6 production observed in E6 expressing cells, cells were treated either with a potent allosteric inhibitor of Akt (Akti), targeting the Akt1 and 2 isoforms [35] or transfected with a plasmid encoding a catalytically inactive Akt mutant (Akt-DN) [36]. Inhibition of Akt by either approach led to significant reduction in *IL-6* mRNA expression (Fig 8C; Akti, p=0.004; Akt-DN, p=0.04 and secretion (Fig 8E; Akti, p= 0.05; Akt-DN, p=0.02).

To confirm that the Akt-mediated increase in IL-6 was transduced via NFκB, we measured p65 phosphorylation levels in C33A cells expression HPV E6 treated with either Akti or co-transfected with Akt-DN. We observed a loss of Akt phosphorylation, indicating that the inhibition strategy was successful, and this was coupled with a reduction in IL-6 protein expression (Fig 8D) and a partial reduction in p65 phosphorylation, suggesting that Akt may act upstream of NFκB in the regulation of IL-6.

We also validated the impact of Akt inhibition on IL-6 production in both HeLa and HPV16+ CaSKi cells. In contrast to HeLa cells, which have wild type *PIK3CA*, CaSKi cells express the E545K mutant of *PIK3CA*, which results in constitutive activation of PI3K/Akt signalling [37]. Interestingly, inhibition of Akt in CaSKi cells had a greater effect in reducing *IL-6* mRNA expression (Fig 8F), IL-6 protein levels (Fig 8G) and secretion (Fig 8H) than in HeLa cells. Furthermore, inhibition of Akt led to a greater reduction in p65 phosphorylation in CaSKi cells. These data confirm that Akt signalling contributes to IL-6 production in HPV cervical cancer cells through the regulation of NFκB.

### IL-6 expression correlates with cervical disease progression

The IL-6/STAT3 signalling axis is deregulated in several cancers [17]. Additionally, IL-6 is over expressed in lung cancer and head and neck cancers [38, 39]. We therefore analysed cervical liquid-based cytology samples from a cohort of HPV16+ patients representing the progression of disease development (CIN1-CIN3) and HPV negative normal cervical tissue to explore its role in cervical disease. Increased levels of *IL-6* mRNA significantly correlated with disease progression through CIN1-CIN3 (Fig 9A; CIN1, p= 0.01; CIN2, p= 0.004; CIN3, p=0.00005), with the greatest increase observed in CIN3 samples when compared with normal cervical tissue. Additionally, IL-6 protein levels also correlated with disease progression (Fig 9B and C; CIN1, p= NS; CIN2, p= 0.0003; CIN3, p=0.04), again showing the largest increase in CIN3. Finally, by mining an available microarray database of normal cervical samples against cervical cancer samples, we observed a statistically significant increase in *IL-6* mRNA expression in the cervical cancer samples (Fig 9D; p=0.03). Together, these data demonstrate that *IL-6* expression is increased in HPV associated cervical disease and in HPV+ cervical cancer.

**Figure 9.**
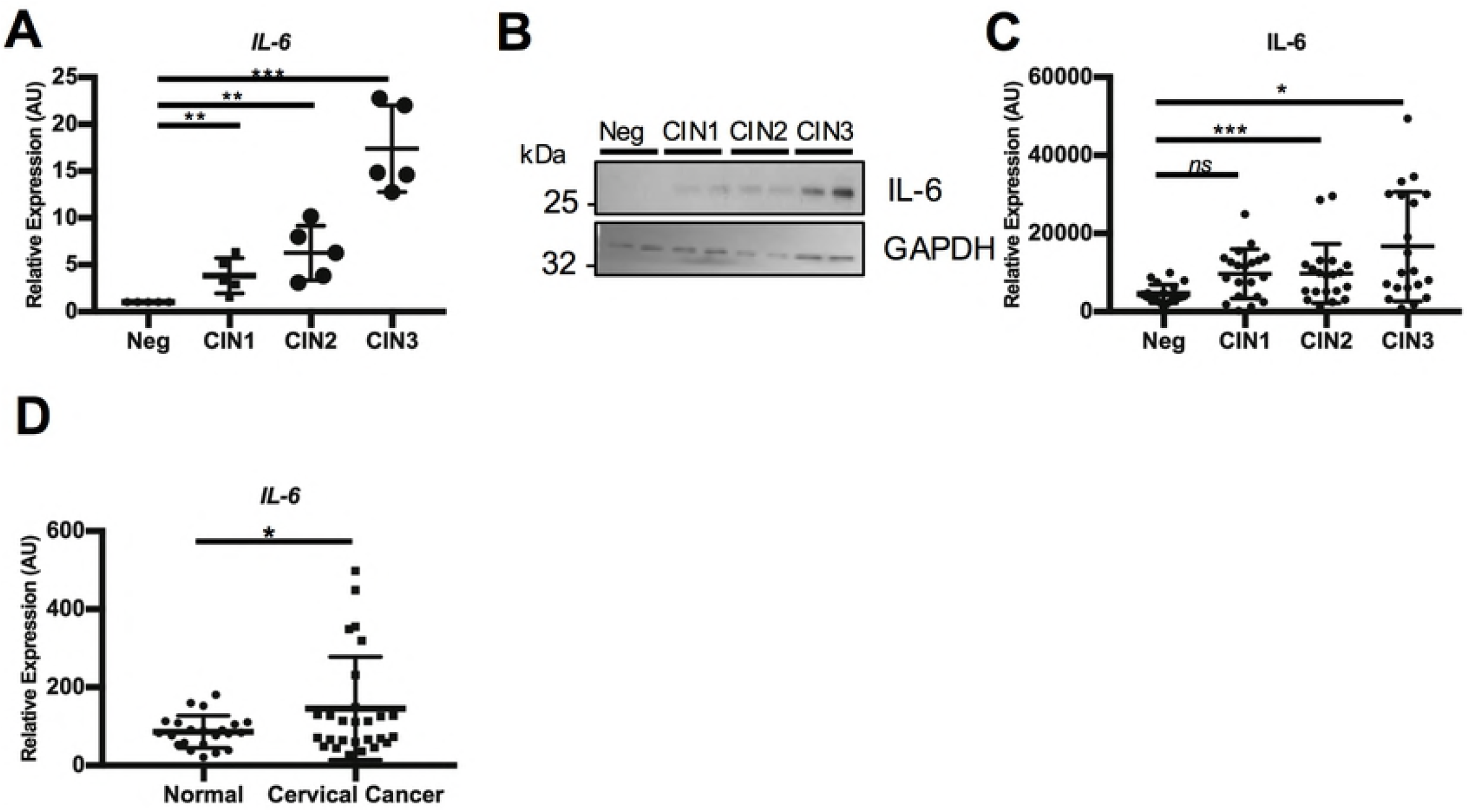
IL-6 expression correlates with cervical disease progression and is up regulated in cervical cancer. **A)** Scatter dot plot of qRT-PCR analysis of RNA extracted from a panel of cytology samples of CIN lesions of increasing grade. Five samples from each clinical grade (neg, CIN I-III) were analysed and mRNA levels were normalized to neg samples. Samples were normalized against U6 mRNA levels. Representative data are presented relative to the HPV- cervical cancer cells. Bars are the means ± standard deviation from at least three biological repeats. *P<0.05, **P<0.01, ***P<0.001 (Student’s t-test). **B)** Representative western blots from cytology samples of CIN lesions of increasing grade analysed IL-6 expression. GAPDH served as a loading control. **C)** Scatter dot plot of densitometry analysis of a panel of cytology samples. Twenty samples from each clinical grade (neg, CIN I-III) were analysed by western blot and densitometry analysis was performed using ImageJ. GAPDH was used as a loading control. **D)** Scatter dot plot of data acquired from the dataset GSE9750 on the GEO database. Arbitrary values for the mRNA expression of IL-6 in normal cervix (n=23) and cervical cancer (n=28) samples were plotted. Error bars represent the mean +/- standard deviation of a minimum of three biological repeats. *P<0.05, **P<0.01, ***P<0.001 (Student’s t-test).

**Figure 10.**
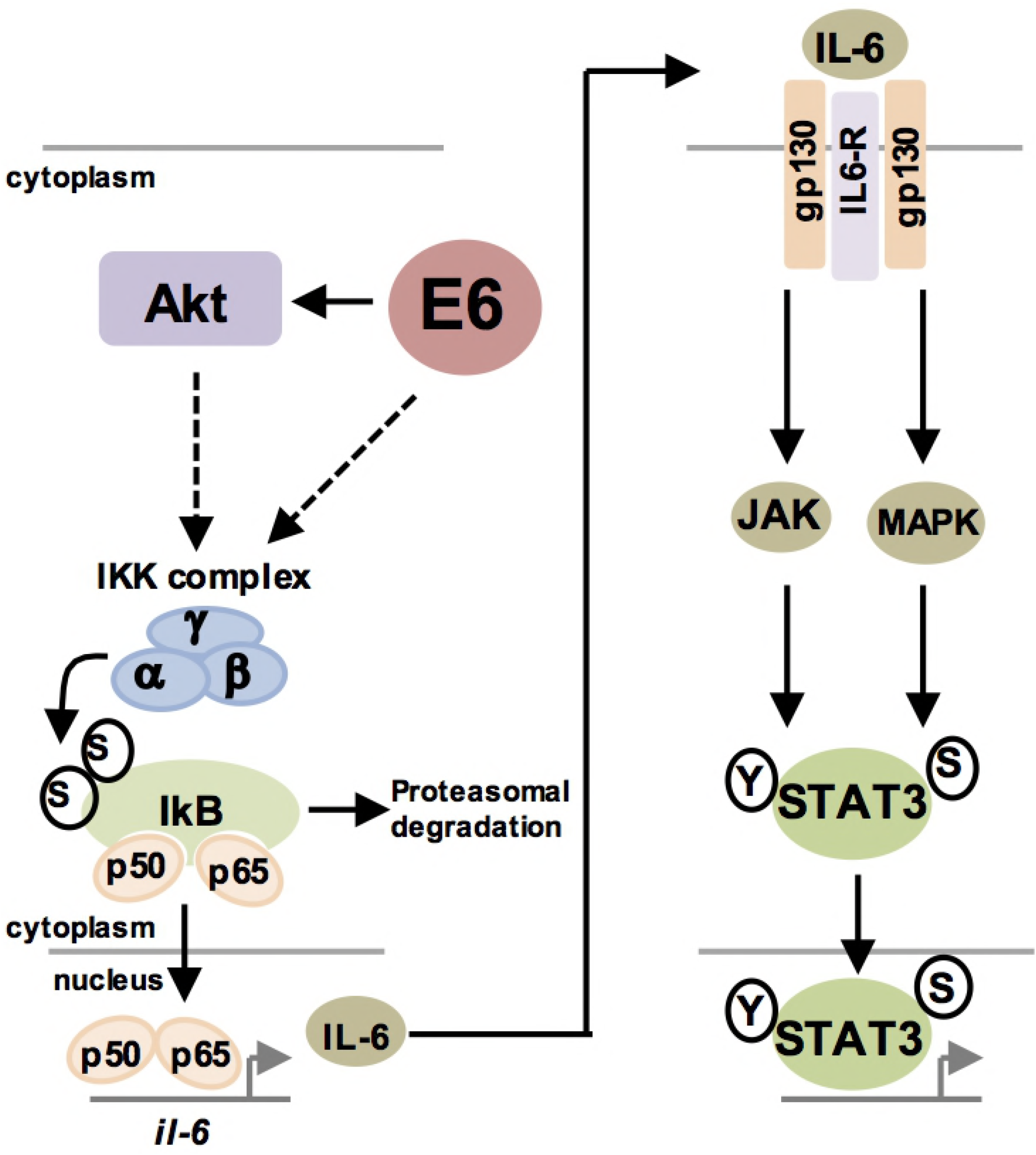
E6 activates an NFκB mediated STAT3 pathway in HPV+ cervical cancer. Schematic diagram of E6 mediated STAT3 signalling in HPV+ cervical cancer cells.

## Discussion

Several oncogenic viruses activate STAT3 to drive cellular proliferation, viral replication and tumourigenesis [18]. Using a primary cell culture model, we have previously shown that E6 activates the STAT3 signalling pathway in primary keratinocytes during the HPV lifecycle [19]. Additionally, we showed that STAT3 activation and expression correlates with cervical disease progression. Although we demonstrated that JAK2 and MAP kinases are responsible for STAT3 phosphorylation in HPV containing cells, the host factors co-opted by HPV E6 required to drive these events are poorly understood. Furthermore, given that previous work has demonstrated that STAT3 inhibition is deleterious to HPV cancer cell survival [20, 21], it was imperative to obtain a more comprehensive understanding of the host pathways necessary for STAT3 activation.

Oncogenic viruses often increase the production of pro-inflammatory cytokines as a conserved mechanism to enhance STAT3 signalling. In particular, IL-6 is over-expressed in diverse cancers and correlates with increased STAT3 activity [17]. IL-6 displays many pleiotropic functions, being both pro-inflammatory and immunosuppressive by interacting with the surrounding stroma of tumours [40]. In HNSCC and oral squamous cell carcinoma, serum levels of IL-6 level are significantly higher than control patients and serum IL-6 is a potential biomarker for these cancers [41]. Additionally, targeting IL-6 in combination with EGFR inhibitors such as Cituximab is currently being investigated as a potential therapy for HNSCC due to the resistance to EGFR inhibition seen in many tumours [42, 43].

IL-6 expression can be regulated by the transcription factor NFκB. In canonical NFκB signalling, various stimuli such as pro-inflammatory cytokines including Tumour Necrosis Factor α (TNFα), initiate a signalling cascade resulting in the phosphorylation of IκBα a negative regulator of the NFκB pathway, by IκB kinases (IKKs). This results in the proteasomal degradation of IκBα and nuclear translocation of the NFκB components p65 and p50 [25].

Kaposi’s Sarcoma associated Herpesvirus (KSHV) down regulates IκBα via the viral miRNA miR-K12-1, leading to enhanced IL-6 expression and STAT3 activation [44]. Both Hepatitis C virus (HCV) core protein and Human Cytomegalovirus (HCMV) US28 protein induce STAT3 phosphorylation and nuclear translocation in an autocrine manner via up regulation of IL-6 [45, 46]

In cervical cancer, IL-6 expression promotes tumour proliferation by inducing vascular epithelial growth factor (VEGF)-dependent angiogenesis in a STAT3 dependent manner [47] and has also been suggested as a potential biomarker [48]. The HPV E6 oncoprotein has been demonstrated to be required for the expression and secretion of IL-6 NSCLC cells [24]; however, the role of E6 in driving IL-6 expression in cervical cancer is unclear. Furthermore, the contribution of IL-6 to STAT3 activation in cervical cancer is poorly defined. Our data show that the increased phosphorylation of STAT3 in HPV+ cervical cancer cells was attributed to an increase in IL-6 expression by HPV E6 and the induction of autocrine/paracrine IL-6/gp130/STAT3 signalling. In cervical cancer cells, EGFR signalling can induce STAT3 activation [49]; however, the data here identified that blockade of IL-6 or gp130 signalling using neutralising antibodies abolished STAT3 phosphorylation, suggesting that IL-6/gp130 is the major determinant for STAT3 phosphorylation in HPV+ cervical cancer cells. We identified NFκB as an essential upstream mediator of IL-6 expression. NFκB is a key component of the inflammatory response, which is a key hallmark of cancer [50]. The induction of inflammation by diverse mechanisms contributes to around 20% of cancers, including the role of inflammatory bowel disease in the development of colorectal cancer (CRC) [51]. Importantly, infection also plays an important role in inflammation-driven cancer; *H. pylori* can lead to the induction of stomach cancers, whilst infection with hepatitis C viruses can lead to development of hepatocellular carcinoma [52-54]. Previous data suggests that inflammation induced by HPV infection may contribute to HPV induced cervical cancers [55, 56]. Indeed, several genes known to be induced by the inflammatory response, including COX-2 [57], are up-regulated in cervical cancer.

The role of NFκB in cervical carcinomas remains controversial, with HPV implicated in both activation and inhibition of the transcription factor [26, 58, 59]. HPV E6 was demonstrated to increase the expression of NFκB components and induce NFκB DNA binding activity, increasing pro-inflammatory cytokine expression [60]. Additionally, E6 reduces the expression of the deubiquitinase CYLD, a known negative regulator of NFκB signalling, in hypoxic cells [27]. In contrast, E6 has been shown to inhibit NFκB transcriptional activity, whilst HPV E7 can attenuate p65 nuclear translocation [61]. The data presented here demonstrates that HPV18 E6 increases the phosphorylation of p65, essential for the nuclear translocation and transactivation ability of p65. Additionally, we demonstrate that NFκB is essential for IL-6 expression in HPV+ cervical cancer cells.

To gain mechanistic insight into the regulation of IL-6 production in HPV+ cervical cancer cells, we focused on the signalling pathways leading to NFκB activation. The protein kinase Akt can regulate NFκB activation under certain circumstances. In PTEN-null cells, Akt activates NFκB through binding of the downstream components mTOR and Raptor to the IKK complex, stimulating NFκB activation [62]. Additionally, Akt can directly phosphorylate and activate IKKα at T23 to enhance p65 phosphorylation [63]. Our data demonstrate that Akt contributes to the phosphorylation of p65 and the expression on IL-6 in HPV+ cervical cancer; however, inhibition of Akt only partially reduced IL-6 expression, suggesting alternative components upstream may be involved in NFκB mediated IL-6 expression.

In cervical cancer, the *PIK3CA* gene is extensively mutated, with the most common mutation (E545K) resulting in constitutive PI3K/Akt signalling [64]. This oncogenic mutation can activate IKK/NFκB signalling and increase IL-6 secretion and paracrine STAT3 activation in epithelial cells [65]. We therefore compared the effect of Akt inhibition on IL-6 production in two cervical cancer cell lines – one with wild type *PIK3CA* (HeLa) and one that has the E545K mutant (CaSKi). We demonstrate that in cells expressing the E545K mutation, and thus having constitutive PI3K/Akt signalling, Akt inhibition has a greater contribution to the NFκB/IL-6 signalling axis than in cells expressing wild type *PIK3CA.* This suggests that targeting the PI3K/Akt pathway in cervical cancers that harbour *PIK3CA* activating mutations, such as E545K, may have therapeutic benefit due to inhibition of NFκB/IL-6 signalling and paracrine STAT3 activation. Indeed, it has recently been demonstrated that the *PIK3CA* E545K mutant confers resistance to cisplatin, suggesting that a combination treatment of cisplatin and a PI3K inhibitor may have synergistic effects in cervical cancer [66].

The data presented here demonstrates that NFκB is essential for the induction of IL-6 and the autocrine/paracrine induction of STAT3 phosphorylation in HPV+ cervical cancer cells. Further, we identify that the protein kinase Akt lies upstream of NFκB/IL-6 signalling as a differential regulator depending on the cellular context due to activating *PIK3CA* mutations. This may allow for the stratification of a subset of cervical cancers that may benefit from a combination treatment of a PI3K inhibitor, such as the pan-PI3K inhibitor buparlisib (BKM120) or the PI3Kα-selective inhibitor alpelisib (BYL719) [67], with standard therapies such as cisplatin.

## Methods and Materials

### Cell Culture

C33A (HPV negative cervical carcinoma), DoTc2 4510 (HPV negative cervical carcinoma), SiHa (HPV 16 positive cervical squamous carcinoma), CaSKi (HPV 16 positive cervical squamous carcinoma), SW756 (HPV 18 positive squamous carcinoma) and HeLa (HPV 18 positive cervical epithelial adenocarcinoma) cells were purchased from ATCC and grown in Dulbecco’s modified Eagle’s media (DMEM) supplemented with 10% Foetal Bovine Serum (FBS) (ThermoFischer Scientific, USA) and 50 U/ml penicillin/streptomycin (Lonza).

### Inhibitors and Cytokines

The IKKα/β inhibitor IKK inhibitor VII was purchased from Calbiochem and used at a final concentration of 5 μM unless otherwise stated. The Akt1/2 inhibitor Akt VIII was purchased from Calbiochem and used at a final concentration of 5 μM unless otherwise stated. Recombinant human IL-6 was purchased from R&D Systems and used at a final concentration of 20 ng/mL unless otherwise stated. Recombinant TNFα was purchased from PeproTech EC Ltd and used at a final concentration of 10 ng/mL. All compounds were used at concentrations required to minimise potential off-target activity. Neutralising IL-6 antibody (ab6628) was purchased from Abcam and used at a final dilution of 1 1:400. Neutralising gp130 antibody (MAB228) was purchased from R&D Systems and used at a final concentration of 1 μg/mL.

### Plasmids and siRNAs

Plasmids expressing HPV18 E6 sequences were amplified from the HPV18 genome and cloned into peGFP-C1 with *SalI* and *XmaI* restriction enzymes. The plasmid driving Firefly luciferase from the IL-6 promoter was a kind gift from Prof Derek Mann (University of Newcastle) and used as previously described [68]; the ConA promoter (that contains tandem NFκB response elements) [69] and a constitutive Renilla luciferase reporter (pRLTK) were previously described [68]. pLNCX1 HA AKT1 were purchased from Addgene (Cambridge, MA, USA) and we thank the principle investigators Jim Darnell, David Baltimore, Linzhao Cheng and Domencino Accili for depositing them. The IκBα S33/36A mutant was a kind gift from Prof Ronald Hay (University of Dundee). For siRNA experiments, two siRNA sequences specifically targeting HPV18 E6 were purchased from GE Healthcare with the following sequences: 5’CUAACACUGGGUUAUACAA’3 and 5’CTAACTAACACTGGGTTAT’3. For HPV16, a single siRNA targeting the HPV16 E6 protein was purchased from Santa Cruz Biotechnology (SCBT; sc-156008). For each experiment, 40nM of pooled siRNA was used and cell lysates were harvested after 72 hours.

### Transfections and mammalian cell lysis

Transient transfections were performed with a DNA to Lipofectamine^®^ 2000 (ThermoFischer) ratio of 1:2.5. 48 h post transfection, cells were lysed in Leeds lysis buffer for western blot [69].

### Western Blotting

Total protein was resolved by SDS-PAGE (10-15% Tris-Glycine), transferred onto Hybond nitrocellulose membrane (Amersham biosciences) and probed with antibodies specific for phospho-STAT3 (S727) (ab32143, abcam), phospho-STAT3 (Y705) (9131, Cell Signalling Technology (CST)), STAT3 (124H6: 9139, CST), phospho-NF-κB p65 (S536) (93H1; 3033, CST), NF-κB p65 (D14E12; 8242, CST), phospho-Akt (T308) (244F9; 4056, CST), phospho-Akt (S473) (D9E; 4060, CST), Akt (9272, CST), IL-6 (ab6672, abcam), HA (HA-7, Sigma H9658), GFP (B-2: sc-9996, SCBT) and GAPDH (G-9, SCBT). Western blots were visualized with species-specific HRP conjugated secondary antibodies (Sigma) and ECL (Thermo/Pierce).

### Retrovirus transduction

pLNCX AKT vector (Addgene, 9006) were transfected into HEK293TT cells with murine retrovirus envelope and GAG/polymerase plasmids (kindly provided by Professor Greg Towers, University College London) using PEI transfection reagent as previously described [19]. After 48 hours the media was removed from the HEK293TT cells and added to HeLa cells for 16 hours. After this time, the virus was removed and replaced with DMEM and cells were harvested 48 hours after transduction.

### Quantitative Real-time PCR

Total RNA was extracted using the E.Z.N.A. Total RNA Kit I (Omega Bio- Tek) according to the manufacture’s protocol. One μg of total RNA was DNase treated following the RQ1 RNase-Free DNase protocol (Promega) and then reverse transcribed with a mixture of random primers and oligo(dT) primers using the qScriptTM cDNA SuperMix (Quanta Biosciences) according to instructions. qRT- PCR was performed using the QuantiFast SYBR Green PCR kit (Qiagen). The PCR reaction was conducted on a Corbett Rotor-Gene 6000 (Qiagen) as follows: initial activation step for 10 min at 95°C and a three-step cycle of denaturation (10 sec at 95°C), annealing (15 sec at 60°C) and extension (20 sec at 72°C) which was repeated 40 times and concluded by melting curve analysis. The data obtained was analysed according to the ΔΔC_t_ method using the Rotor-Gene 6000 software [70]. Specific primers were used for each gene analysed and are shown in Table 1. U6 served as normaliser gene.

### Luciferase Reporter Assays

Cells were seeded into 12 well dishes and transfected the following day using PEI with reporter plasmids expressing firefly luciferase under the control of the *IL-6* promoter or the *ConA* promoter, which contains tandem repeats of a κB-response element [68, 71]. Where appropriate, cells were co-transfected with plasmids expressing GFP or GFP-E6. To normalise for transfection efficiency, pRLTK Renilla luciferase reporter plasmid was added to each transfection. After 24 hours, samples were lysed in passive lysis buffer (Promega) and activity measured using a dual-luciferase reporter assay system (Promega) as described [72].

### Immunofluorescent Staining

Cells were seeded onto coverslips and, 24 hr later, were transfected as required. 24 hr after transfection, cells were fixed with 4 % paraformaldehyde for 10 min and then permeabilised with 0.1 % (v/v) Triton for 15 minutes. Cells were then incubated in primary antibodies in PBS with 4 % BSA overnight at 4°C. Primary antibodies were used at a concentration of 1:400. Cells were washed thoroughly in PBS and then incubated with Alex-fluor conjugated secondary antibodies 594 and Alexa 488 (1:1000) (Invitrogen) in PBS with 4% BSA for 2 hours. DAPI was used to visualise nuclei. Coverslips were mounted onto slides with Prolong Gold (Invitrogen).

### ELISA

The human IL-6 DuoSet^®^ ELISA was purchased from R&D Systems and was used according to the manufacturer’s instructions.

### Microarray analysis

For microarray analysis, a dataset of 28 cervical cancer cases and 23 normal cervix samples was utilised. Microarray data was obtained from GEO database accession number GSE9750.

### Statistical analysis

Where indicated, data was analysed using a two-tailed, unpaired Student’s t-test. Figure Legends

**Supplementary Figure 1. HPV16 E6 induces IL-6 expression**. **A)** CaSKi cells were transfected with HPV16 E6 specific siRNA and analysed for IL-6 mRNA expression by qRT-PCR. Samples were normalized against U6 mRNA levels. **B)** Representative western blot of CaSKi cells transfected with a pool of two specific siRNAs against HPV16 E6 and analysed for the expression of IL-6. Knockdown of HPV16 E6 was confirmed using an antibody against HPV16 E6 and p53. GAPDH served as a loading control. **G)** CaSKi cells were transfected with a pool of two specific siRNAs against HPV16 E6. The culture medium was analysed for IL-6 protein by ELISA. Data are representative of at least three biological independent repeats. Error bars represent the mean +/- standard deviation of a minimum of three biological repeats. *P<0.05, **P<0.01, ***P<0.001 (Student’s t-test).

**Supplementary Figure 2. NF-κB inhibition does not affect exogenous IL-6 mediated STAT3 signalling**. **A)** C33A were treated with DMSO or IKKi or transfected with pcDNA or IκBm before treatment with 20 ng/mL recombinant human IL-6 for 30 mins. Cell lysates were analysed for the phosphorylated and total forms of p65 and STAT3. GAPDH served as a loading control. Data are representative of at least three biological independent repeats. **B)** C33A were treated with DMSO or IKKi or transfected with pcDNA or IκBm before treatment with 20 ng/mL recombinant human IL-6 for 30 mins. Cells were analysed by immunofluorescence staining for total STAT3 (green) and counterstained with DAPI to highlight the nuclei (blue in the merged panels). Scale bar, 20 μm.

## Acknowledgements

We are grateful to William Sellers for providing the retroviral Akt expression vectors through the Addgene repository. We thank the Scottish HPV Investigators Network (SHINE) and Sheila Graham (University of Glasgow) for providing HPV positive biopsy samples. Martin Stacey (University of Leeds) provided help and reagents for ELISA experiments, Derek Mann (University of Newcastle) and Ron Hay (University of Dundee) provided plasmid reagents. Stephen Griffin (University of Leeds) and Matthew Reeves (University College London) provided helpful discussions.

## Author contributions

Conceived and designed the experiments: ELM and AM. Performed the experiments: ELM. Wrote the manuscript: ELM and AM. Critically analysed the manuscript: ELM and AM.

## References

1. Hausen zur H. Papillomaviruses and cancer: from basic studies to clinical application. Nat Rev Cancer. 2002;2: 342–350. doi:10.1038/nrc798

2. Crosbie EJ, Einstein MH, Franceschi S, Kitchener HC. Human papillomavirus and cervical cancer. Lancet. 2013;382: 889–899. doi:10.1016/S0140-6736(13)60022-7

3. Wasson CW, Morgan EL, Müller M, Ross RL, Hartley M, Roberts S, et al. Human papillomavirus type 18 E5 oncogene supports cell cycle progression and impairs epithelial differentiation by modulating growth factor receptor signalling during the virus life cycle. Oncotarget. Impact Journals LLC; 2017;8: 103581–103600. doi:10.18632/oncotarget.21658

4. Zhang B, Srirangam A, Potter DA, Roman A. HPV16 E5 protein disrupts the c-Cbl-EGFR interaction and EGFR ubiquitination in human foreskin keratinocytes. Oncogene. Nature Publishing Group; 2005;24: 2585–2588. doi:10.1038/sj.onc.1208453

5. Shostak K, Zhang X, Hubert P, Göktuna SI, Jiang Z, Klevernic I, et al. NF-κB-induced KIAA1199 promotes survival through EGFR signalling. Nat Commun. Nature Publishing Group; 2014;5: 5232. doi:10.1038/ncomms6232

6. Hu T, Li C. Convergence between Wnt-β-catenin and EGFR signaling in cancer. Mol Cancer. BioMed Central; 2010;9: 236. doi:10.1186/1476-4598-9-236

7. Bello JOM, Nieva LO, Paredes AC, Gonzalez AMF, Zavaleta LR, Lizano M. Regulation of the Wnt/β-Catenin Signaling Pathway by Human Papillomavirus E6 and E7 Oncoproteins. Viruses. Multidisciplinary Digital Publishing Institute; 2015;7: 4734–4755. doi:10.3390/v7082842

8. He C, Mao D, Hua G, Lv X, Chen X, Angeletti PC, et al. The Hippo/YAP pathway interacts with EGFR signaling and HPV oncoproteins to regulate cervical cancer progression. EMBO Mol Med. EMBO Press; 2015;7: 1426–1449. doi:10.15252/emmm.201404976

9. Rodríguez MI, Finbow ME, Alonso A. Binding of human papillomavirus 16 E5 to the 16 kDa subunit c (proteolipid) of the vacuolar H+-ATPase can be dissociated from the E5-mediated epidermal growth factor receptor overactivation. Oncogene. Nature Publishing Group; 2000;19: 3727–3732. doi:10.1038/sj.onc.1203718

10. Wetherill LF, Holmes KK, Verow M, Müller M, Howell G, Harris M, et al. High-risk human papillomavirus E5 oncoprotein displays channel-forming activity sensitive to small-molecule inhibitors. J Virol. American Society for Microbiology; 2012;86: 5341–5351. doi:10.1128/JVI.06243-11

11. Wetherill LF, Wasson CW, Swinscoe G, Kealy D, Foster R, Griffin S, et al. Alkyl-imino sugars inhibit the pro-oncogenic ion channel function of human papillomavirus (HPV) E5. Antiviral Res. 2018;158: 113–121. doi:10.1016/j.antiviral.2018.08.005

12. Scheffner M, Huibregtse JM, Vierstra RD, Howley PM. The HPV-16 E6 and E6-AP complex functions as a ubiquitin-protein ligase in the ubiquitination of p53. Cell. 1993;75: 495–505.

13. Banerjee NS, Wang H-K, Broker TR, Chow LT. Human papillomavirus (HPV) E7 induces prolonged G2 following S phase reentry in differentiated human keratinocytes. J Biol Chem. 2011;286: 15473–15482. doi:10.1074/jbc.M110.197574

14. Moody CA, Laimins LA. Human papillomaviruses activate the ATM DNA damage pathway for viral genome amplification upon differentiation. Galloway D, editor. PLoS Pathog. Public Library of Science; 2009;5: e1000605. doi:10.1371/journal.ppat.1000605

15. Yu H, Lee H, Herrmann A, Buettner R, Jove R. Revisiting STAT3 signalling in cancer: new and unexpected biological functions. Nat Rev Cancer. 2014;14: 736–746. doi:10.1038/nrc3818

16. Carpenter RL, Lo H-W. STAT3 Target Genes Relevant to Human Cancers. Cancers (Basel). Multidisciplinary Digital Publishing Institute; 2014;6: 897–925. doi:10.3390/cancers6020897

17. Johnson DE, O’Keefe RA, Grandis JR. Targeting the IL-6/JAK/STAT3 signalling axis in cancer. Nat Rev Clin Oncol. Nature Publishing Group; 2018;15: 234–248. doi:10.1038/nrclinonc.2018.8

18. Roca Suarez AA, Van Renne N, Baumert TF, Lupberger J. Viral manipulation of STAT3: Evade, exploit, and injure. Hobman TC, editor. PLoS Pathog. 2018;14: e1006839. doi:10.1371/journal.ppat.1006839

19. Morgan EL, Wasson CW, Hanson L, Kealy D, Pentland I, McGuire V, et al. STAT3 activation by E6 is essential for the differentiation-dependent HPV18 life cycle. Galloway DA, editor. PLoS Pathog. 2018;14: e1006975. doi:10.1371/journal.ppat.1006975

20. Shukla S, Mahata S, Shishodia G, Pandey A, Tyagi A, Vishnoi K, et al. Functional regulatory role of STAT3 in HPV16-mediated cervical carcinogenesis. Williams BO, editor. PLoS ONE. Public Library of Science; 2013;8: e67849. doi:10.1371/journal.pone.0067849

21. Shukla S, Shishodia G, Mahata S, Hedau S, Pandey A, Bhambhani S, et al. Aberrant expression and constitutive activation of STAT3 in cervical carcinogenesis: implications in high-risk human papillomavirus infection. Mol Cancer. BioMed Central; 2010;9: 282. doi:10.1186/1476-4598-9-282

22. Silver JS, Hunter CA. gp130 at the nexus of inflammation, autoimmunity, and cancer. J Leukoc Biol. 2010;88: 1145–1156. doi:10.1189/jlb.0410217

23. Yu H, Pardoll D, Jove R. STATs in cancer inflammation and immunity: a leading role for STAT3. Nat Rev Cancer. 2009;9: 798–809. doi:10.1038/nrc2734

24. Cheng Y-W, Lee H, Shiau M-Y, Wu T-C, Huang T-T, Chang Y-H. Human papillomavirus type 16/18 up-regulates the expression of interleukin-6 and antiapoptotic Mcl-1 in non-small cell lung cancer. Clin Cancer Res. 2008;14: 4705–4712. doi:10.1158/1078-0432.CCR-07-4675

25. Hinz M, Scheidereit C. The IκB kinase complex in NF-κB regulation and beyond. EMBO Rep. 2014;15: 46–61. doi:10.1002/embr.201337983

26. James MA, Lee JH, Klingelhutz AJ. Human papillomavirus type 16 E6 activates NF-kappaB, induces cIAP-2 expression, and protects against apoptosis in a PDZ binding motif-dependent manner. J Virol. 2006;80: 5301–5307. doi:10.1128/JVI.01942-05

27. An J, Mo D, Liu H, Veena MS, Srivatsan ES, Massoumi R, et al. Inactivation of the CYLD deubiquitinase by HPV E6 mediates hypoxia-induced NF-kappaB activation. Cancer Cell. 2008;14: 394–407. doi:10.1016/j.ccr.2008.10.007

28. Kroll M, Margottin F, Kohl A, Renard P, Durand H, Concordet JP, et al. Inducible degradation of IkappaBalpha by the proteasome requires interaction with the F-box protein h-betaTrCP. J Biol Chem. 1999;274: 7941–7945.

29. Zhang Q, Lenardo MJ, Baltimore D. 30 Years of NF-κB: A Blossoming of Relevance to Human Pathobiology. Cell. 2017;168: 37–57. doi:10.1016/j.cell.2016.12.012

30. Litwin TR, Clarke MA, Dean M, Wentzensen N. Somatic Host Cell Alterations in HPV Carcinogenesis. Viruses. Multidisciplinary Digital Publishing Institute; 2017;9: 206. doi:10.3390/v9080206

31. Hussain AR, Ahmed SO, Ahmed M, Khan OS, Abdulmohsen Al S, Platanias LC, et al. Cross-talk between NFκB and the PI3-kinase/AKT pathway can be targeted in primary effusion lymphoma (PEL) cell lines for efficient apoptosis. Gallyas F, editor. PLoS ONE. Public Library of Science; 2012;7: e39945. doi:10.1371/journal.pone.0039945

32. Ahmad A, Biersack B, Li Y, Kong D, Bao B, Schobert R, et al. Targeted regulation of PI3K/Akt/mTOR/NF-κB signaling by indole compounds and their derivatives: mechanistic details and biological implications for cancer therapy. Anticancer Agents Med Chem. NIH Public Access; 2013;13: 1002–1013.

33. Cahill CM, Rogers JT. Interleukin (IL) 1beta induction of IL-6 is mediated by a novel phosphatidylinositol 3-kinase-dependent AKT/IkappaB kinase alpha pathway targeting activator protein-1. J Biol Chem. 2008;283: 25900–25912. doi:10.1074/jbc.M707692200

34. Spangle JM, Münger K. The human papillomavirus type 16 E6 oncoprotein activates mTORC1 signaling and increases protein synthesis. J Virol. 2010;84: 9398–9407. doi:10.1128/JVI.00974-10

35. Zhao Z, Leister WH, Robinson RG, Barnett SF, Defeo-Jones D, Jones RE, et al. Discovery of 2,3,5-trisubstituted pyridine derivatives as potent Akt1 and Akt2 dual inhibitors. Bioorg Med Chem Lett. 2005;15: 905–909. doi:10.1016/j.bmcl.2004.12.062

36. Street A, Macdonald A, McCormick C, Harris M. Hepatitis C virus NS5A-mediated activation of phosphoinositide 3-kinase results in stabilization of cellular beta-catenin and stimulation of beta-catenin-responsive transcription. J Virol. American Society for Microbiology Journals; 2005;79: 5006–5016. doi:10.1128/JVI.79.8.5006-5016.2005

37. Kang S, Bader AG, Vogt PK. Phosphatidylinositol 3-kinase mutations identified in human cancer are oncogenic. Proc Natl Acad Sci USA. 2005;102: 802–807. doi:10.1073/pnas.0408864102

38. Yadav A, Kumar B, Datta J, Teknos TN, Kumar P. IL-6 promotes head and neck tumor metastasis by inducing epithelial-mesenchymal transition via the JAK-STAT3-SNAIL signaling pathway. Mol Cancer Res. 2011;9: 1658–1667. doi:10.1158/1541-7786.MCR-11-0271

39. Sriuranpong V, Park JI, Amornphimoltham P, Patel V, Nelkin BD, Gutkind JS. Epidermal growth factor receptor-independent constitutive activation of STAT3 in head and neck squamous cell carcinoma is mediated by the autocrine/paracrine stimulation of the interleukin 6/gp130 cytokine system. Cancer Res. 2003;63: 2948–2956.

40. Fisher DT, Appenheimer MM, Evans SS. The two faces of IL-6 in the tumor microenvironment. Semin Immunol. 2014;26: 38–47. doi:10.1016/j.smim.2014.01.008

41. Gao J, Zhao S, Halstensen TS. Increased interleukin-6 expression is associated with poor prognosis and acquired cisplatin resistance in head and neck squamous cell carcinoma. Oncol Rep. Spandidos Publications; 2016;35: 3265–3274. doi:10.3892/or.2016.4765

42. Stanam A, Love-Homan L, Joseph TS, Espinosa-Cotton M, Simons AL. Upregulated interleukin-6 expression contributes to erlotinib resistance in head and neck squamous cell carcinoma. Mol Oncol. 2015;9: 1371–1383. doi:10.1016/j.molonc.2015.03.008

43. Stabile LP, Egloff AM, Gibson MK, Gooding WE, Ohr J, Zhou P, et al. IL6 is associated with response to dasatinib and cetuximab: Phase II clinical trial with mechanistic correlatives in cetuximab-resistant head and neck cancer. Oral Oncol. 2017;69: 38–45. doi:10.1016/j.oraloncology.2017.03.011

44. Chen M, Sun F, Han L, Qu Z. Kaposi’s sarcoma herpesvirus (KSHV) microRNA K12-1 functions as an oncogene by activating NF-κB/IL-6/STAT3 signaling. Oncotarget. Impact Journals; 2016;7: 33363–33373. doi:10.18632/oncotarget.9221

45. Tacke RS, Tosello-Trampont A, Nguyen V, Mullins DW, Hahn YS. Extracellular hepatitis C virus core protein activates STAT3 in human monocytes/macrophages/dendritic cells via an IL-6 autocrine pathway. J Biol Chem. American Society for Biochemistry and Molecular Biology; 2011;286: 10847–10855. doi:10.1074/jbc.M110.217653

46. Slinger E, Maussang D, Schreiber A, Siderius M, Rahbar A, Fraile-Ramos A, et al. HCMV-encoded chemokine receptor US28 mediates proliferative signaling through the IL-6-STAT3 axis. Sci Signal. 2010;3: ra58–ra58. doi:10.1126/scisignal.2001180

47. Wei L-H, Kuo M-L, Chen C-A, Chou C-H, Lai K-B, Lee C-N, et al. Interleukin-6 promotes cervical tumor growth by VEGF-dependent angiogenesis via a STAT3 pathway. Oncogene. Nature Publishing Group; 2003;22: 1517–1527. doi:10.1038/sj.onc.1206226

48. Song Z, Lin Y, Ye X, Feng C, Lu Y, Yang G, et al. Expression of IL-1α and IL-6 is Associated with Progression and Prognosis of Human Cervical Cancer. Med Sci Monit. International Scientific Information, Inc; 2016;22: 4475–4481. doi:10.12659/MSM.898569

49. Knudsen SLJ, Mac ASW, Henriksen L, van Deurs B, Grøvdal LM. EGFR signaling patterns are regulated by its different ligands. Growth Factors. Taylor & Francis; 2014;32: 155–163. doi:10.3109/08977194.2014.952410

50. Karin M. NF-kappaB as a critical link between inflammation and cancer. Cold Spring Harb Perspect Biol. 2009;1: a000141–a000141. doi:10.1101/cshperspect.a000141

51. Janakiram NB, Rao CV. The role of inflammation in colon cancer. Adv Exp Med Biol. Basel: Springer Basel; 2014;816: 25–52. doi:10.1007/978-3-0348-0837-8_2

52. Polk DB, Peek RM. Helicobacter pylori: gastric cancer and beyond. Nat Rev Cancer. Nature Publishing Group; 2010;10: 403–414. doi:10.1038/nrc2857

53. Zhang X-Y, Zhang P-Y, Aboul-Soud MAM. From inflammation to gastric cancer: Role of Helicobacter pylori. Oncol Lett. Spandidos Publications; 2017;13: 543–548. doi:10.3892/ol.2016.5506

54. Castello G, Scala S, Palmieri G, Curley SA, Izzo F. HCV-related hepatocellular carcinoma: From chronic inflammation to cancer. Clin Immunol. 2010;134: 237–250. doi:10.1016/j.clim.2009.10.007

55. Castle PE, Hillier SL, Rabe LK, Hildesheim A, Herrero R, Bratti MC, et al. An association of cervical inflammation with high-grade cervical neoplasia in women infected with oncogenic human papillomavirus (HPV). Cancer Epidemiol Biomarkers Prev. 2001;10: 1021–1027.

56. Fernandes JV, DE Medeiros Fernandes TAA, DE Azevedo JCV, Cobucci RNO, DE Carvalho MGF, Andrade VS, et al. Link between chronic inflammation and human papillomavirus-induced carcinogenesis (Review). Oncol Lett. 2015;9: 1015–1026. doi:10.3892/ol.2015.2884

57. Subbaramaiah K, Dannenberg AJ. Cyclooxygenase-2 transcription is regulated by human papillomavirus 16 E6 and E7 oncoproteins: evidence of a corepressor/coactivator exchange. Cancer Res. 2007;67: 3976–3985. doi:10.1158/0008-5472.CAN-06-4273

58. Hasan U. Human papillomavirus (HPV) deregulation of Toll-like receptor 9. Oncoimmunology. Taylor & Francis; 2014;3: e27257. doi:10.4161/onci.27257

59. Ma W, Tummers B, van Esch EMG, Goedemans R, Melief CJM, Meyers C, et al. Human Papillomavirus Downregulates the Expression of IFITM1 and RIPK3 to Escape from IFNγ- and TNFα-Mediated Antiproliferative Effects and Necroptosis. Front Immunol. 2016;7: 496. doi:10.3389/fimmu.2016.00496

60. Nees M, Geoghegan JM, Hyman T, Frank S, Miller L, Woodworth CD. Papillomavirus type 16 oncogenes downregulate expression of interferon-responsive genes and upregulate proliferation-associated and NF-kappaB-responsive genes in cervical keratinocytes. J Virol. 2001;75: 4283–4296. doi:10.1128/JVI.75.9.4283-4296.2001

61. Vandermark ER, Deluca KA, Gardner CR, Marker DF, Schreiner CN, Strickland DA, et al. Human papillomavirus type 16 E6 and E 7 proteins alter NF-κB in cultured cervical epithelial cells and inhibition of NF-κB promotes cell growth and immortalization. Virology. 2012;425: 53–60. doi:10.1016/j.virol.2011.12.023

62. Dan HC, Cooper MJ, Cogswell PC, Duncan JA, Ting JP-Y, Baldwin AS. Akt-dependent regulation of NF-{kappa}B is controlled by mTOR and Raptor in association with IKK. Genes Dev. Cold Spring Harbor Lab; 2008;22: 1490–1500. doi:10.1101/gad.1662308

63. Bai D, Ueno L, Vogt PK. Akt-mediated regulation of NFkappaB and the essentialness of NFkappaB for the oncogenicity of PI3K and Akt. Int J Cancer. John Wiley & Sons, Ltd; 2009;125: 2863–2870. doi:10.1002/ijc.24748

64. Bader AG, Kang S, Vogt PK. Cancer-specific mutations in PIK3CA are oncogenic in vivo. Proc Natl Acad Sci USA. 2006;103: 1475–1479. doi:10.1073/pnas.0510857103

65. Hutti JE, Pfefferle AD, Russell SC, Sircar M, Perou CM, Baldwin AS. Oncogenic PI3K mutations lead to NF-κB-dependent cytokine expression following growth factor deprivation. Cancer Res. American Association for Cancer Research; 2012;72: 3260–3269. doi:10.1158/0008-5472.CAN-11-4141

66. Arjumand W, Merry CD, Wang C, Saba E, McIntyre JB, Fang S, et al. Phosphatidyl inositol-3 kinase (PIK3CA) E545K mutation confers cisplatin resistance and a migratory phenotype in cervical cancer cells. Oncotarget. 2016;7: 82424–82439. doi:10.18632/oncotarget.10955

67. Massacesi C, Di Tomaso E, Urban P, Germa C, Quadt C, Trandafir L, et al. PI3K inhibitors as new cancer therapeutics: implications for clinical trial design. Onco Targets Ther. Dove Press; 2016;9: 203–210. doi:10.2147/OTT.S89967

68. Macdonald A, Crowder K, Street A, McCormick C, Saksela K, Harris M. The hepatitis C virus non-structural NS5A protein inhibits activating protein-1 function by perturbing ras-ERK pathway signaling. J Biol Chem. 2003;278: 17775–17784. doi:10.1074/jbc.M210900200

69. Richards KH, Wasson CW, Watherston O, Doble R, Blair GE, Wittmann M, et al. The human papillomavirus (HPV) E7 protein antagonises an Imiquimod-induced inflammatory pathway in primary human keratinocytes. Sci Rep. Nature Publishing Group; 2015;5: 12922. doi:10.1038/srep12922

70. Livak KJ, Schmittgen TD. Analysis of relative gene expression data using real-time quantitative PCR and the 2(-Delta Delta C(T)) Method. Methods. 2001;25: 402–408. doi:10.1006/meth.2001.1262

71. Griffiths DA, Abdul-Sada H, Knight LM, Jackson BR, Richards K, Prescott EL, et al. Merkel cell polyomavirus small T antigen targets the NEMO adaptor protein to disrupt inflammatory signaling. J Virol. 2013;87: 13853–13867. doi:10.1128/JVI.02159-13

72. Macdonald A, Mazaleyrat S, McCormick C, Street A, Burgoyne NJ, Jackson RM, et al. Further studies on hepatitis C virus NS5A-SH3 domain interactions: identification of residues critical for binding and implications for viral RNA replication and modulation of cell signalling. J Gen Virol. Microbiology Society; 2005;86: 1035–1044. doi:10.1099/vir.0.80734-0

